# Selective Immune Silencing by Targeted TGF-β Mimics

**DOI:** 10.64898/2026.01.19.700410

**Authors:** Qinli Sun, Masato Ogishi, Alison K. Barrett, Hua Jiang, Hao Yan, Elsa Sola, Jie Zhang, Peng Xiao, Huiyun Lyu, Ahmad Salehi, Honghui Liu, Qizhi Tang, Mark M. Davis, Robert S. Negrin, Tobias V. Lanz, K. Christopher Garcia

## Abstract

Depletion of pathogenic T and B cells is a pillar of therapies for autoimmune, inflammatory, and transplantation-related immunological diseases. However, adverse events, safety concerns in immunocompromised patients, and disease relapse limit clinical utility. Here, we exploit the immunosuppressive properties of a transforming growth factor beta (TGF-β) mimic repurposed from helminths for cell type–specific therapeutic “silencing”, as a new approach to complement existing therapies. Mouse CD4⁺ and CD8⁺ T cell–targeted TGF-β agonists selectively and potently silence antigen-specific T cell responses in OVA-immunized mice by suppressing pro-inflammatory effector, cytotoxic, and T follicular helper programs, while skewing cells toward a quiescent state biased toward regulatory and type 17 T cell phenotypes. Similarly, human CD4⁺ and CD8⁺ T cell–targeted TGF-β agonists precisely and effectively suppress live-attenuated influenza vaccine (LAIV)-induced T cell activation and expansion in human spleen organoids. Correspondingly, CD4⁺ T cell-targeted TGF-β agonist effectively ameliorated disease activity and promoted disease remission in CD4⁺ T cell-driven models of autoimmune neuroinflammation and allergic airway inflammation, demonstrating efficacy in both prophylactic and established inflammatory settings. Moreover, both CD4⁺ and CD8⁺ T cell-targeted TGF-β agonists ameliorated disease activity in graft-versus-host disease. Additionally, a human CD19⁺ B cell–targeted TGF-β agonist robustly inhibits germinal center B cell-to-plasmablast maturation and antibody responses in LAIV-stimulated human spleen organoids. These early-stage results suggest that cell-selective TGF-β agonism merits further investigation as a versatile therapeutic approach for the precise silencing of pathogenic adaptive immune responses.

## Introduction

Dysregulated adaptive T and B lymphocytes drive immunopathology across a broad spectrum of immune-mediated diseases, including inflammatory, allergic, autoimmune, and transplant-related disorders^1,2^. Therapeutic strategies aimed at depleting or functionally restraining pathogenic T and B cells have been extensively explored and, in many cases, transformative^3–5^. T cell–suppressing agents such as tacrolimus are widely used to prevent graft rejection in hematopoietic stem cell and solid organ transplantation^6,7^. In contrast, CD3-specific antibodies, including teplizumab and otelixizumab, modulate T cell function and induce transient peripheral T cell reduction, with clinical activity reported in settings such as type 1 diabetes and transplantation^8,9^. B cell–depleting monoclonal antibodies (mAbs), including the CD20-targeting agents ocrelizumab and obinutuzumab, have demonstrated therapeutic efficacy in multiple sclerosis^10,11^ and active lupus nephritis^12^. More recently, CD19-targeted CAR T cell therapy has shown encouraging clinical efficacy in refractory systemic lupus erythematosus and other autoimmune diseases^13,14^.

While valuable, lymphocyte-depleting therapies have important limitations that merit development of alternative or complementary approaches. First, the loss of adaptive immune cells increases the susceptibility to infection. For example, broad T cell–targeted therapies such as Teplizumab induce T-lymphopenia and cause viral reactivation^15^. Strategies that selectively suppress pathogenic T-cell types while preserving other protective immunity would therefore be preferable but are not yet available. Long-term B cell depletion also causes hypogammaglobulinemia and an increased risk of serious infections, although this risk can be mitigated with intravenous immunoglobulin therapy^16^. Second, T- and B-cell–directed depletion strategies, including anti-CD3 antibodies and CD19-targeted CAR T cell therapies, can trigger inflammatory toxicities, most notably cytokine release syndrome (CRS), and immune effector cell-associated neurotoxicity syndrome (ICANS)^17,18^. These adverse events frequently necessitate additional immunosuppression and treatment interruption. Lastly, although disease relapses following CD19-targeted CAR-T therapy have rarely been reported across multiple autoimmune diseases^14^, long-term relapse risk remains under investigation. Importantly, given the longer life expectancy of autoimmune patients compared to cancer patients, disease relapse would necessitate repeated CAR-T cycles, imposing substantial economic and quality-of-life burdens. Therefore, immunosuppression strategies that are more cell-type-specific and mechanistically orthogonal to existing cytolytic therapeutics could enhance the flexibility of clinical management for autoimmune diseases.

Transforming growth factor-β (TGF-β) is an essential immunoregulatory cytokine that mediates immune tolerance and suppresses inflammation^19^. TGF-β acts as a critical suppressor of T cell activation and expansion and plays a pivotal role in regulating the differentiation of distinct T cell subsets^20^; for example, it promotes peripherally induced regulatory CD4^+^ T cell (pTreg) differentiation^21^, drives type 17 helper CD4⁺ T (Th17) cell development^22,23^, and supports tissue-resident memory T cell formation^24^. TGF-β has also been reported to suppress B cell expansion and hyperresponsiveness while regulating antibody class-switch recombination^25^. Importantly, impaired TGF-β signaling *in vivo* has been associated with heightened inflammation and autoimmunity^26–28^, highlighting its potential as a promising immunosuppressive therapeutic target. However, poor drug-like properties, latent homodimer formation, and pro-fibrotic or tumorigenic effects in non-immune cells have severely limited its safe and effective *in vivo* use^29^. Moreover, broad targeting may lead to a “sink effect,” reducing its availability and limiting its ability to adequately suppress pathogenic immune cell populations. Therefore, given these limitations, the therapeutic potential of TGF-β has been only modestly explored^30^; however, a more precise strategy for targeting TGF-β agonism to specific cell types or tissue locations could substantially enhance its clinical utility.

To explore the concept of selective immune silencing via TGF-β signaling, we focused on a helminth-derived TGF-β mimic 1 (TGM1)^31,32^, a well-established TGF-β agonist that was recently shown to robustly induce antigen-specific pTreg differentiation *in vivo* in combination with IL-2^33^. We fused the low-affinity domains of TGM1 to high-affinity T cell– or B cell–specific targeting modules, enabling selective activation of TGF-β signaling in T and B cells, particularly antigen-activated clones, thereby achieving precise immunosuppression while minimizing off-target effects. We show that CD4⁺ and CD8⁺ T cell–targeted, as well as CD19^+^ B cell–targeted, TGF-β agonists selectively suppress activation and expansion of their respective target cell types in *in vivo* mouse models and human organoids, thereby effectively attenuating T cell–driven autoimmune neuroinflammation, airway allergy, and graft-versus-host disease (GVHD) in prophylactic settings and, in some models, under established inflammatory conditions, as well as suppressing B cell–derived influenza-specific antibody responses. Mechanistically, gain-of-function treatment with these TGF-β agonists reveals that strong TGF-β signaling skews naïve T cells toward a “quiescent” state by blocking inflammatory and T follicular helper (Tfh) differentiation pathways and antagonizing type I interferon signaling. It also potently suppresses the differentiation of antibody-producing B cells. Overall, these findings highlight a promising therapeutic strategy for pathogenic T- and B-cell–driven diseases through selective TGF-β agonism.

## Results

### Design and characterization of mouse T cell-targeted TGF-β agonists

Full-length helminth TGM1 (TGM1FL), the TGF-β agonist used here, comprises five domains: domains 1–3 engage TGF-β receptors with low affinity, whereas domains 4–5 bind co-receptor CD44 with high affinity, enabling broad immunosuppression and facilitating immune evasion in infected hosts^32,34^. To generate CD4⁺ and CD8⁺ T cell–targeted TGF-β agonists in mice (**Fig. 1A**), we replaced the CD44-binding domains of TGM1FL with either a high-affinity mouse CD4-specific single-chain variable fragment (scFv)^35^ (TGM1-mCD4) or a mouse CD8α-specific variable domain of heavy-chain-only antibody (V_H_H)^36^ (TGM1-mCD8), linked via a flexible GSG linker (**Fig. 1B and S1A)**. This design redirects TGF-β signaling mediated by the low-affinity TGM1 domains 1–3 (hereafter TGM1, to distinguish it from the five-domain parent protein TGM1FL) selectively to CD4⁺ or CD8α⁺ T cells, while minimizing engagement of non-target cells (**Fig. 1A**). Unless stated otherwise, low-affinity TGM1 domains 1–3 is the untargeted control used throughout this study. We further fused the constructs to mouse serum albumin (MSA) to extend *in vivo* half-life (**Fig. 1B and S1A)**. The resulting proteins were expressed at high yield in Expi293 cells and behaved as monodisperse species with favorable biochemical properties (**Fig. S1B and S1C)**. Phosphorylated SMAD2/3 (pSMAD2/3) assays in mCD4- or mCD8α-transduced TGF-β reporter HEK-293 cells showed that TGM1-mCD4 and TGM1-mCD8 left-shifted the pSMAD2/3 dose–response curves by ∼4 and ∼3 orders of magnitude, respectively, compared with low-affinity TGM1 alone, whereas the targeting modules alone had no detectable activity (**Fig. 1C and S1D)**. Moreover, pSMAD2/3 assays in untransduced HEK293 cells, mouse NIH/3T3 fibroblasts, and bEnd.3 endothelial cells showed that, whereas recombinant human TGF-β1 (hTGFβ1) efficiently activated SMAD2/3 at low concentrations, neither low-affinity TGM1 nor the targeted TGM1s elicited detectable SMAD2/3 activation at low concentrations, with activity becoming detectable only at concentrations of 100 nM or higher (**Fig. S1E)**. Together, these results demonstrate that high-affinity CD4 or CD8α targeting substantially enhances functional TGF-β receptor engagement by the low-affinity TGM1 signaling core, enabling potent TGF-β signaling selectively in the respective target cell populations while maintaining minimal activity in non-target cells and thereby creating a broad safety window relative to the widespread activity of native TGF-β across TGF-β receptor-expressing cell types.

**Figure 1.**
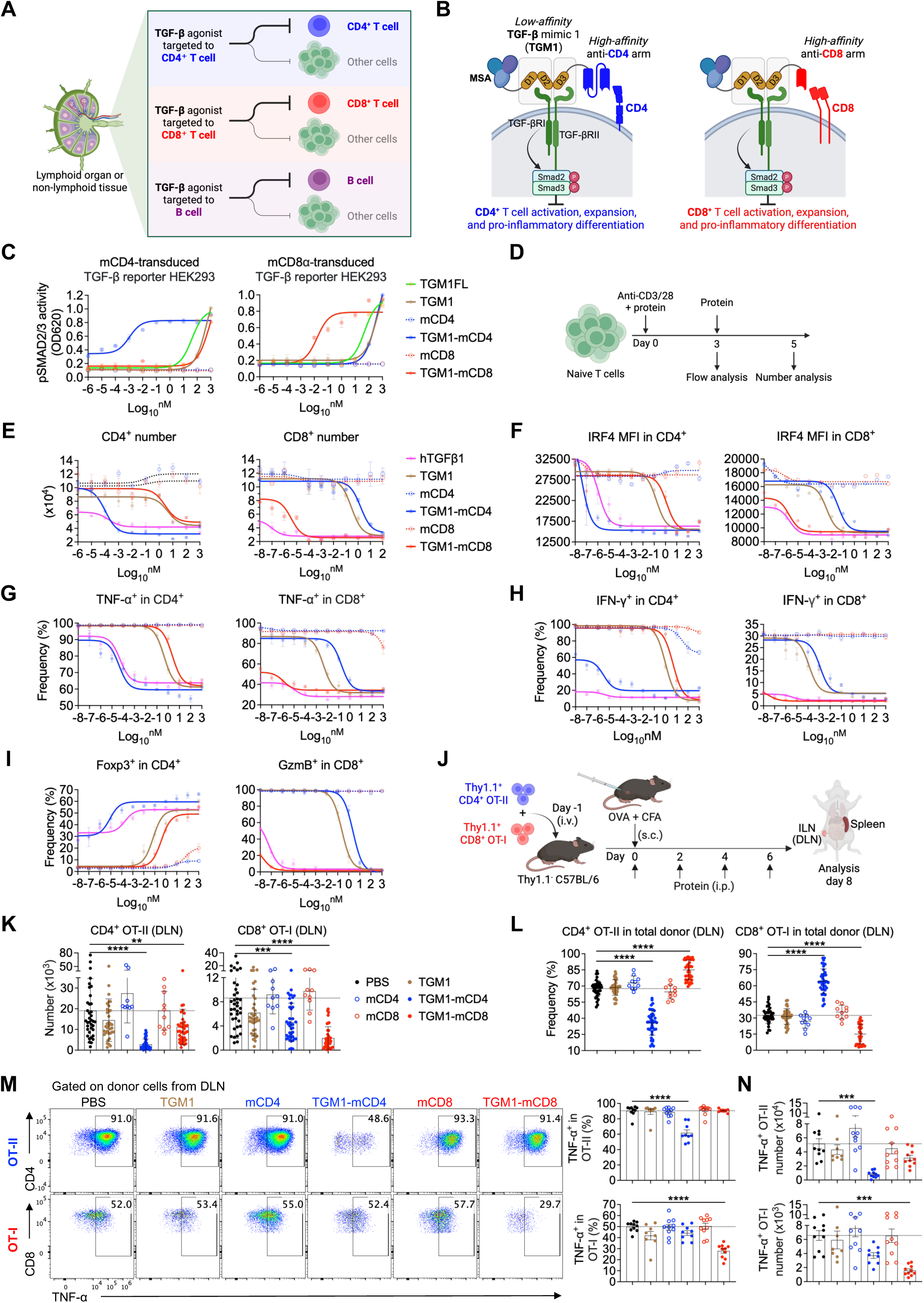
Design and characterization of mouse T cell-targeted TGF-β agonists. **A.** Schematic diagram illustrating selective *in vivo* suppression of T or B cells by targeted TGF-β agonists. **B.** Schematic diagram illustrating the molecular design of T cell–targeted TGF-β agonists. **C.** Dose–response curves showing pSMAD2/3 levels in mouse CD4⁺- or CD8α-transduced TGF-β reporter HEK293 cells using a pSMAD-induced secreted alkaline phosphatase (SEAP) assay. **D.** Experimental design of *in vitro* mouse T cell culture in the presence of the indicated proteins. **E.** Dose–response curves showing numbers of *in vitro* cultured CD4⁺ or CD8⁺ T cells treated with the indicated proteins on day 5. **F.** Dose–response curves showing IRF4 expression levels in *in vitro* cultured CD4⁺ or CD8⁺ T cells. **G.** Dose–response curves showing TNF-α production in *in vitro* cultured CD4⁺ or CD8⁺ T cells. **H.** Dose–response curves showing IFN-γ production in *in vitro* cultured CD4⁺ or CD8⁺ T cells. **I.** Dose–response curves showing Foxp3 and Granzyme B expression in *in vitro* cultured CD4⁺ and CD8⁺ T cells, respectively. **J.** Schematic diagram illustrating the experimental design of naïve OT-II and OT-I cell transfer, OVA immunization, and protein administration. **K.** Quantification of donor Thy1.1^+^ OT-II and OT-I cell numbers in DLNs from the indicated mice (n = 40 mice per group). **L.** Quantification of the proportions of Thy1.1^+^ OT-II or OT-I cells among total donor Thy1.1^+^ cells (OT-II + OT-I) in DLNs from the indicated mice (n = 40 mice per group). **M.** Representative flow cytometry plots and quantification of TNF-α–producing OT-II or OT-I cell frequencies in DLNs from the indicated mice (n = 10 mice per group). **N.** Quantification of TNF-α–producing OT-II or OT-I cell numbers in DLNs from the indicated mice. Data in **C**, **E-I, M and N** are representative of two or three independent experiments. Data in **K and L** are pooled from four independent experiments. Data in **C, E-I, M, and N** are presented as mean ± s.e.m. Data in **K and L** are presented as mean ± s.d. The statistics were obtained by one-way ANOVA coupled with Dunnett’s multiple-comparisons test **(K-N)**.

To validate selective T cell suppression, naïve CD4⁺ or CD8⁺ T cells were cultured *in vitro* and activated with anti-CD3/CD28 in the presence of the indicated proteins (**Fig. 1D**). Prior to this, to assess whether the targeting arms interfere with binding of commercial anti-mCD4 and anti-mCD8α flow cytometry antibodies, we incubated isolated primary mouse CD4⁺ or CD8⁺ T cells with the corresponding proteins at 37 °C for 1 hour prior to staining. The data showed that the targeting arms partially competed with binding of certain antibody clones (**Fig. S1F and S1G)**. Accordingly, we selected non-competing clones (anti-mCD4, clone RM4-4; anti-mCD8α, clone 5H10-1) for subsequent gating of the corresponding mouse CD4⁺ or CD8⁺ T cells (**Fig. S1F and S1G)**. Functionally, similar to hTGFβ1, TGM1-mCD4 and TGM1-mCD8 potently and selectively suppressed T cell receptor (TCR)-driven expansion of CD4⁺ and CD8⁺ T cells, respectively, shifting the dose–inhibition curves for cell number by ∼5–6 orders of magnitude relative to TGM1 alone or the alternative T cell–targeted construct (**Fig. 1E**). Additionally, TGM1-mCD4 and TGM1-mCD8 effectively suppressed their respective T cell activation, as shown by leftward shifts in dose–inhibition curves for CD25, CD69, GITR, IRF4, and T-bet expression, as well as for inflammatory cytokines IFN-γ and TNF-α, in a manner similar to hTGFβ1 (**Fig. 1F-H, S2A, and S2B)**. Moreover, consistent with hTGFβ1, TGM1-mCD4 effectively promoted Foxp3⁺ Treg induction in CD4⁺ T cells, whereas TGM1-mCD8 effectively suppressed Granzyme B production in CD8⁺ T cells (**Fig. 1I**). Together, these data demonstrate that CD4⁺ or CD8⁺ T cell–targeted TGM1 selectively and effectively suppresses TCR-driven activation, inflammatory cytokine production, and expansion of the corresponding T cell subset *in vitro*.

### Mouse T cell–targeted TGF-β agonists selectively and potently suppress antigen-specific T cell responses in OVA-immunized mice

Next, we assessed the *in vivo* safety and efficacy of TGM1-mCD4 and TGM1-mCD8 in regulating antigen-specific T cell responses. Naïve Thy1.1⁺ CD4⁺ OT-II and CD8⁺ OT-I T cells were co-transferred into Thy1.1⁻ recipient mice, followed by immunization with OVA emulsified in complete Freund’s adjuvant (CFA) to induce a robust inflammatory response (**Fig. 1J**). To determine the protein dose for *in vivo* experiments, we referred to the dosing regimen used in our previous study^33^ and performed an *in vivo* dose-titration experiment (10, 30, 70, and 100 pmol per dose, administered every other day starting on day 0) to evaluate donor OT-II and OT-I cell responses in draining lymph nodes (DLNs) (**Fig. 1J and S2C)**. TGM1-mCD4 and TGM1-mCD8 selectively suppressed TNF-α production in their respective target populations, CD4⁺ OT-II and CD8⁺ OT-I cells, with detectable activity at 30 pmol and progressively greater effects at 70 and 100 pmol (**Fig. S2C)**. In contrast, low-affinity TGM1 alone and the mCD4 and mCD8 targeting antibodies alone showed minimal effects even at 100 pmol (**Fig. S2C)**, demonstrating the target-dependent activity of TGM1-mCD4 and TGM1-mCD8 and indicating that substantial biological activity can be achieved at relatively low doses. Based on these results, we selected approximately 40–50 pmol per dose (TGM1, 3.82–4.78 μg; TGM1-mCD4, 4.91–6.14 μg; TGM1-mCD8, 4.43–5.54 μg; mCD4, 3.81–4.76 μg; and mCD8, 3.25–4.06 μg) for subsequent *in vivo* studies to achieve robust target-cell activity while limiting systemic exposure and potential off-target TGF-β signaling.

Furthermore, *in vivo* studies using the selected protein dose showed that body weight and spleen weight were comparable between mice treated with TGM1-mCD4, TGM1-mCD8, or the targeting arms alone, and PBS-treated controls (**Fig. S3A-C)**, indicating favorable *in vivo* safety and preservation of immune homeostasis. Analysis of donor OT-II and OT-I cells in DLNs and spleens showed that, consistent with their *in vitro* selectivity, TGM1-mCD4 markedly reduced OT-II cell numbers in both DLNs and spleens, whereas TGM1-mCD8 selectively and robustly reduced OT-I cell numbers (**Fig. 1K, S3D, and S3E)**. Notably, TGM1-mCD4 decreased the proportion of OT-II cells and correspondingly increased the frequency of OT-I cells among total donor cells (OT-II + OT-I), while TGM1-mCD8 produced the opposite effect (**Fig. 1L and S3F)**, highlighting their selective suppression of the targeted T cell subsets. In contrast, the mCD4 and mCD8 targeting arms alone, as well as low-affinity TGM1, showed minimal effects on donor T cell populations (**Fig. 1K, 1L, S3E, and S3F)**. Moreover, both TGM1-mCD4 and TGM1-mCD8 significantly reduced the frequency and absolute number of TNF-α–producing cells within their respective antigen-specific donor cell populations in both DLNs and spleens (**Fig. 1M, 1N, and S3G)**, indicating potent and selective inhibition of inflammatory differentiation. Given that CD8α is also expressed on classical dendritic cells (cDCs)^37^, we further examined the effect of TGM1-mCD8 on this population (**Fig. S3H)**. The data showed that TGM1-mCD8 had no substantial impact on CD8α⁺ cDC abundance or on CD80 and CD86 expression in this population ∼48 hours after OVA and CFA immunization (**Fig. S3I and S3J)**, suggesting that the observed *in vivo* effects are likely primarily attributable to direct suppression of T cells. Together, these results demonstrate that TGM1-mCD4 and TGM1-mCD8 selectively and potently suppress OVA-specific CD4⁺ and CD8⁺ T cell responses, consistent with previous studies of T cell–intrinsic TGF-β receptor deficiency^38^.

### Phenotypic characterization of antigen-stimulated T and B cells *in vivo* following treatment with T cell–targeted TGF-β agonists

We next analyzed the effects of T cell-targeted TGF-β agonists on antigen-specific T cell lineage differentiation in OVA-immunized mice. TGM1-mCD4 increased the frequencies of Foxp3⁺ Treg cells (∼5%) and Foxp3⁻ RORγt⁺ cells (∼3%) in the DLNs, compared with PBS controls (**Fig. 2A, 2B, and (S4A)**. Although TGM1-mCD4 substantially increased the Foxp3⁺ population, reaching a frequency of ∼5% and representing an approximately 10-fold increase over PBS controls, the magnitude of Treg induction was markedly lower than that observed in our previous study using a targeted TGF-β–IL-2 co-agonist, which induced Foxp3⁺ Treg cells at frequencies of 60–80% among antigen-specific CD4⁺ T cells^33^. TGM1-mCD4 also increased the frequency of IL-17A-producing Th17 cells (∼2%) among OT-II cells in the DLNs (**Fig. 2A, 2B, and S4A)**. Importantly, TGM1-mCD4 markedly suppressed T follicular helper (Tfh) cell development, as evidenced by a pronounced reduction in PD-1⁺Bcl6⁺ cells in both donor OT-II cells and endogenous CD44⁺CD4⁺ T cells in the DLNs (**Fig. 2A, 2B, S4A, and S4B)**. Consistently, germinal center B (GC B) cell differentiation was severely impaired in DLNs of TGM1-mCD4–treated mice, as shown by significantly reduced frequencies and numbers of Fas⁺GL7⁺ B cells (**Fig. 2C, 2D, and S4C)**, accompanied by markedly decreased serum OVA-specific IgG1 levels, compared with PBS controls (**Fig. 2E**). Quantification of the numbers of each CD4⁺ T cell subset further revealed a marked reduction in both Tfh and IFN-γ–producing Th1 cells, whereas Treg and Th17 cell numbers remained largely unchanged (**Fig. 2B, S4A, and S4B)**. In contrast, neither untargeted TGM1 nor TGM1–mCD8 had appreciable effects on the differentiation or abundance of donor OT-II cell subsets in the DLNs (**Fig. 2A, 2B, S4A, and S4B)**. Together, these data indicate that TGM1-mCD4 preferentially suppresses inflammatory Tfh and Th1 differentiation with limited lineage skewing toward Treg or Th17 cells, thereby leading to potent inhibition of antigen-specific humoral immune responses.

**Figure 2.**
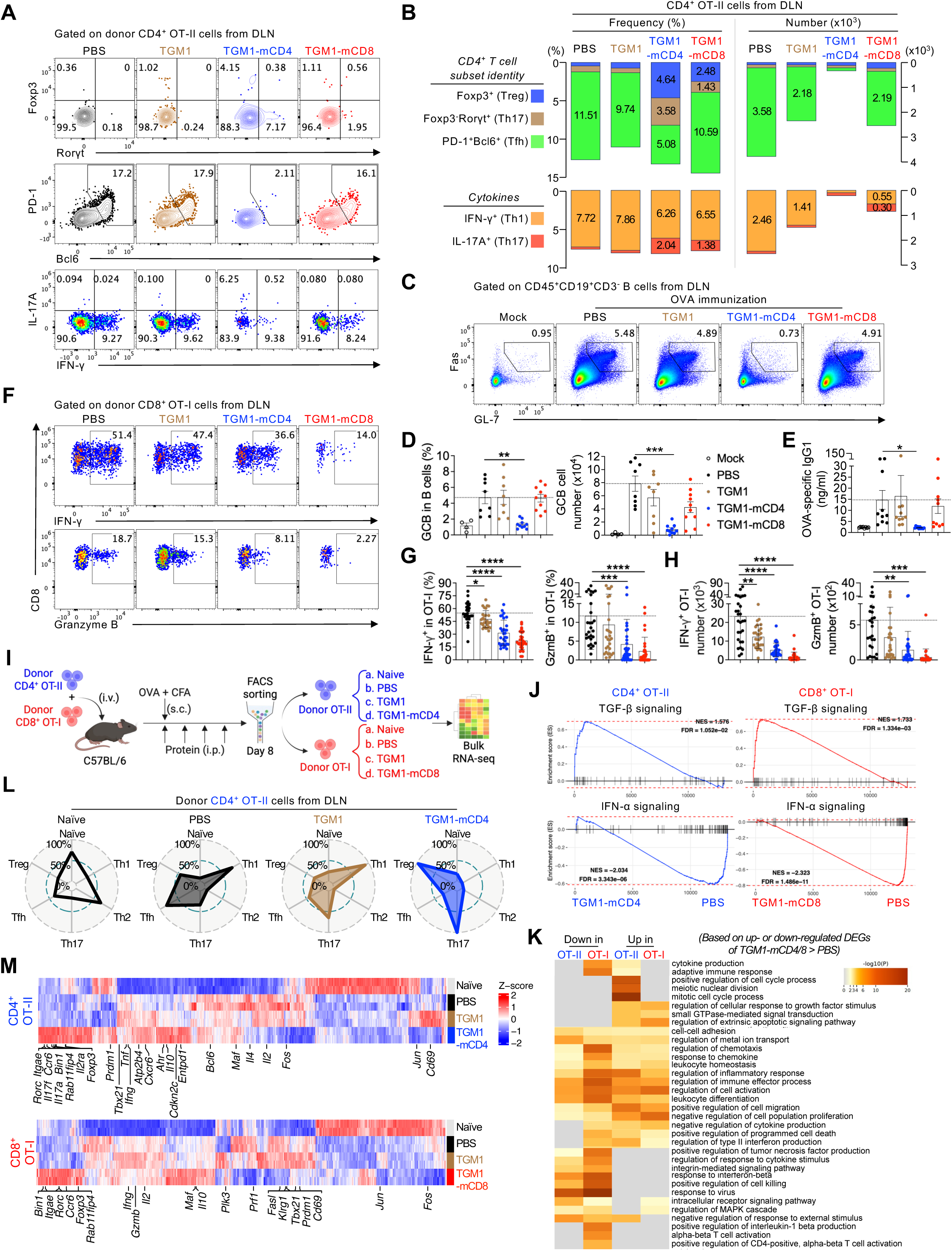
Mouse T cell-targeted TGF-β agonists silence antigen-specific T and B cell responses in OVA-immunized mice. **A.** Representative flow cytometry plots showing Foxp3, RORγt, PD-1, and Bcl6 expression, as well as IL-17A and IFN-γ production in OT-II cells from DLNs of the indicated mice (n = 20 mice per group). **B.** Bar plots showing the frequencies and numbers of the indicated CD4⁺ T cell populations within OT-II cells from DLNs of the indicated mice (n = 20 mice per group). **C.** Representative flow cytometry plots showing Fas⁺GL-7⁺ GC B cell frequencies and numbers in DLNs of the indicated mice (Mock, n = 4 mice; PBS, n = 8 mice; TGM1, n = 8 mice; TGM1-mCD4, n = 10 mice; TGM1-mCD8, n = 10 mice per group). **D.** Quantification of Fas⁺GL-7⁺ GC B cell frequencies and numbers in DLNs of the indicated mice. **E.** Quantification of serum OVA-specific IgG1 levels in the indicated mice. **F.** Representative flow cytometry plots showing IFN-γ– and Granzyme B–producing cell frequencies in OT-II or OT-I cells from DLNs of the indicated mice (n = 20 mice per group). **G.** Quantification of IFN-γ– and Granzyme B–producing cell frequencies in OT-II or OT-I cells from DLNs of the indicated mice. **H.** Quantification of IFN-γ– and Granzyme B–producing OT-II and OT-I cell numbers in DLNs of the indicated mice. **I.** Schematic diagram illustrating the experimental design for OT-II and OT-I bulk RNA-seq sample preparation. **J.** GSEA showing enrichment of TGF-β and IFN-α signaling signatures in OT-II or OT-I cells from targeted TGM1 groups versus PBS. **K.** Gene Ontology (GO) analysis showing representative biological pathways enriched among up- or down-regulated DEGs in OT-II or OT-I cells from targeted TGM1 versus PBS groups. **L.** Spider plots showing enrichment scores of various CD4⁺ T cell subset signatures in OT-II cell transcriptomes from the indicated groups. **M.** Heatmaps showing expression levels of DEGs in OT-II or OT-I cells between indicated groups, with representative feature genes highlighted. Data in **A, B, and F-H** are pooled from two independent experiments. Data in **C-E** are representative of two independent experiments. Data in **D and E** are presented as mean ± s.e.m. Data in **G and H** are presented as mean ± s.d. The statistics were obtained by one-way ANOVA coupled with Dunnett’s multiple-comparisons test **(D, E, G, and H)**.

Consistently, TGM1-mCD8 markedly reduced both the frequency and number of IFN-γ– and granzyme B–producing OT-I cells in the DLNs (**Fig. 2F–H**), whereas TGM1-mCD4 also led to a modest reduction, likely secondary to impaired CD4⁺ T cell help for CD8⁺ T cell responses^39^. These results indicate that CD8⁺ T cell–targeted TGF-β agonists potently suppress cytotoxic effector CD8⁺ T cell differentiation *in vivo*.

To gain insight into how T cell–targeted TGF-β agonists regulate antigen-specific T cell transcriptomes, we isolated donor OT-II and OT-I cells from the DLNs of the OVA-immunized mice described above, together with naïve OT-II and OT-I cells, for bulk RNA sequencing (RNA-seq) (**Fig. 2I**). Principal component analysis (PCA) and gene clustering analysis revealed that OT-II and OT-I cells treated with their corresponding targeted TGM1 constructs adopted transcriptional profiles distinct from PBS controls and shifted away from the naïve state, indicating induction of a unique cellular program (**Fig. S5A and S5B)**. In contrast, OT-II and OT-I cells treated with untargeted TGM1 clustered closely with PBS controls in PCA space, underscoring its minimal activity in the absence of T cell–targeting modules (**Fig. S5A and S5B).** Gene set enrichment analysis (GSEA) and gene expression heatmaps further showed that OT-II and OT-I cells treated with their corresponding targeted TGM1 constructs exhibited enrichment of TGF-β signaling gene signatures relative to PBS controls (**Fig. 2J and S5C)**, indicating that the observed transcriptional changes were likely driven by active TGF-β signaling delivered by the targeted TGM1 agonists. Notably, GSEA, gene expression heatmaps, and pathway enrichment analyses revealed that OT-II and OT-I cells treated with targeted TGM1 constructs exhibited significantly reduced type I interferon (IFN-α) signaling signatures compared with PBS controls at the transcriptional level (**Fig. 2J, 2K, and S5D)**, suggesting suppression of inflammatory pathways by targeted TGF-β agonists.

Next, we further analyzed the altered T cell lineage programs induced by targeted TGM1 constructs at the transcriptional level. Consistent with the protein-level data, GSEA showed that TGM1-mCD4 skewed OT-II cells toward Treg and Th17 transcriptional programs, with marked suppression of Th1 and Tfh signatures (**Fig. 2L**), accompanied by increased expression of Treg and Th17-associated genes (Treg: *Foxp3, Il10, Il2ra, and Entpd1*; Th17: *Rorc, Ahr, Maf, Il17a, Il17f,* and *Ccr6*) and reduced expression of Th1 and Tfh-associated genes (e.g., *Fos, Il2*, and *Bcl6*), as shown in the gene expression heatmap, compared with PBS controls (**Fig. 2M**). Moreover, TGM1-mCD8 substantially reduced the expression of cytotoxic and effector function–related genes (e.g., *Tbx21, Prdm1, Gzmb, Prf1, Fasl,* and *Klrg1*) compared with PBS controls, suggesting suppression of cytotoxic CD8⁺ T cell differentiation (**Fig. 2M**). Notably, we also observed increased expression of Treg- and Th17-associated genes (e.g., *Rorc, Foxp3, Ccr6,* and *Il10*) at transcriptional level in OT-I cells (**Fig. 2M**), indicating that targeted TGM1 induces partially overlapping transcriptional changes in both CD4⁺ and CD8⁺ T cells. This was further supported by substantial overlap in differentially expressed genes (DEGs) between OT-II and OT-I cells (**Fig. S5E)**.

Additionally, upregulated DEGs shared by OT-II and OT-I cells treated with targeted TGM1 versus PBS included genes associated with T cell migration, tissue homing, and tissue retention, such as *Itgae, Gpr15, Ccr4, Ccr6, Ccr8, and Cx3cr1* **(Fig. S5F)**. In contrast, downregulated DEGs included members of the GIMAP family (*Gimap3*, *Gimap4*, and *Gimap5*), which are involved in T cell survival and homeostasis, as well as integrin family members (*Itga4, Itgb1, Itgb3*, and *Itga7*), which regulate cell adhesion, migration, and interactions with the tissue microenvironment **(Fig. S5F)**. Together, these transcriptional changes suggest that, in addition to regulating T cell differentiation, targeted TGM1 may modulate gene programs associated with T cell migration, tissue homing, adhesion, and retention.

Collectively, these data indicate that T cell–targeted TGM1 programs T cells into a “quiescent” transcriptional state *in vivo* characterized by suppression of type I effector, cytotoxic, and Tfh programs, a shift toward Treg and Th17-associated signatures, and concomitant downregulation of type I interferon pathways.

### Human T cell–targeted TGF-β agonists potently suppress antigen-stimulated T cell responses in human spleen organoids

Given that TGM1 shows cross-reactivity with both mouse and human TGF-β receptors^31^, we next generated human CD4⁺ and CD8⁺ T cell–targeted TGF-β agonists by similarly fusing the low-affinity TGM1 domains 1–3 with either a high-affinity human CD4-targeting nanobody (Nb)^40^ (TGM1-hCD4) or a human CD8α-targeting V_H_H^41^ (TGM1-hCD8), respectively (**Fig. S6A and S6B)**. pSMAD assays in human CD4- or CD8α-transduced TGF-β reporter HEK293 cells showed that both TGM1-hCD4 and TGM1-hCD8 shifted the pSMAD2/3 dose–stimulation curve leftward by approximately 4 and 8 orders of magnitude, respectively, compared with low-affinity TGM1 alone (**Fig. 3A and S6C)**, demonstrating markedly enhanced functional potency conferred by the targeting domains. Flow cytometry confirmed that the hCD4-targeting arm does not compete with commercial anti-CD4 antibodies (clone SK3, clone RPA-T4, and clone A161A1) (**Fig. S6D)**, and that the hCD8α-targeting arm does not compete with a commercial anti-CD8α antibody (clone S21010A) (**Fig. S6E)**; these clones were therefore selected for use in subsequent staining experiments. We then cultured CD4⁺ or CD8⁺ T cells isolated from peripheral blood mononuclear cells (PBMCs) of healthy donors *in vitro* under anti-CD3/CD28 stimulation in the presence of these proteins **(Fig. S6F)**. As observed with hTGFβ1, TGM1–hCD4 and TGM1–hCD8 each markedly and selectively reduced CD4⁺ and CD8⁺ T cell numbers, respectively, in a dose-dependent manner (**Fig. 3B**). In contrast, low-affinity TGM1, the targeting arms alone, and the alternative targeted constructs had no detectable effect (**Fig. 3B**). Moreover, consistent with hTGFβ1 activity, TGM1-hCD4 significantly increased the proportion of FOXP3⁺ CD4⁺ T cells, whereas TGM1-hCD8 strongly reduced the frequency of Granzyme B–producing CD8⁺ T cells in a dose-responsive manner **(Fig. S6G)**. Together, these results demonstrate that human CD4⁺- and CD8α-targeted TGM1 selectively and effectively modulate their respective human T cell subsets *in vitro*.

**Figure 3.**
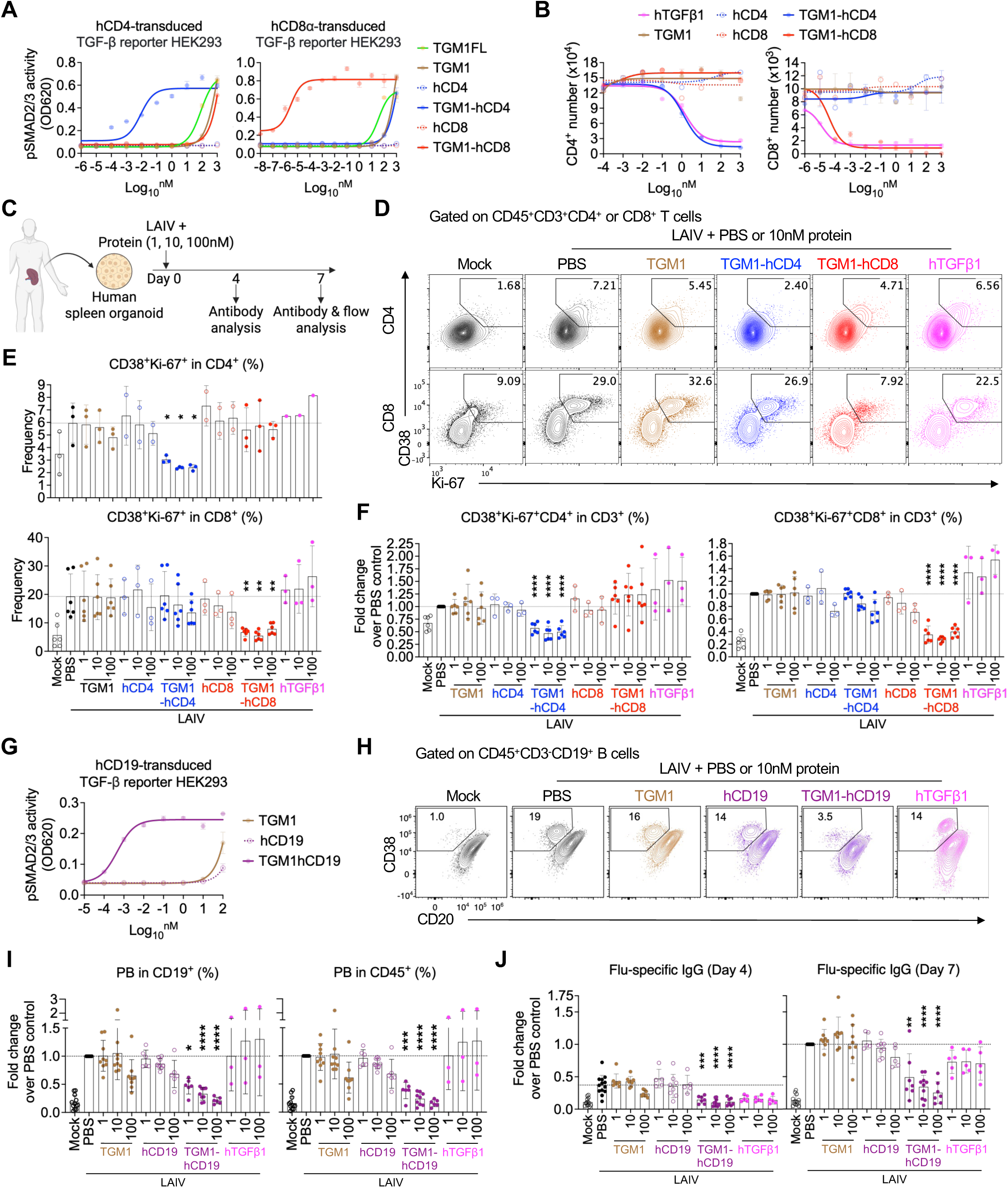
Human T and B cell-targeted TGF-β agonists silence LAIV-induced T and B cell responses in human spleen organoids. **A.** Dose–response curves showing pSMAD2/3 levels in human CD4- or CD8α-transduced TGF-β reporter HEK293 cells measured using a pSMAD-induced SEAP assay. **B.** Dose–response curves showing numbers of *in vitro*–cultured human CD4⁺ or CD8⁺ T cells treated with the indicated proteins on day 5. **C.** Schematic diagram illustrating the experimental design of human spleen organoid culture, LAIV stimulation, and protein treatment. **D.** Representative flow cytometry plots showing CD38⁺Ki-67⁺ cell frequencies in CD4⁺ or CD8⁺ T cells from the indicated groups. “Mock” refers to human spleen organoids cultured in vitro without LAIV stimulation, whereas “PBS” refers to LAIV-stimulated human spleen organoids treated with PBS as the vehicle control in the absence of the indicated proteins. **E.** Quantification of CD38⁺Ki-67⁺ cell frequencies in CD4⁺ or CD8⁺ T cells from the indicated groups. **F.** Quantification of CD38⁺Ki-67⁺ CD4⁺ or CD8⁺ T cells among total CD3⁺ T cells from the indicated groups. **G.** Dose–response curves showing pSMAD2/3 levels in human CD19-transduced TGF-β reporter HEK293 cells measured using a pSMAD-induced SEAP assay. **H.** Representative flow cytometry plots showing plasmablast (PB) frequencies among B cells from the indicated groups. **I.** Quantification of plasmablast frequencies among B cells and total CD45⁺ leukocytes from the indicated groups. **J.** Quantification of influenza (flu)-specific IgG levels in culture supernatants from the indicated groups on days 4 and 7. Data in **A, B, and G** are representative of two independent experiments. Data in **D-F and H-J** are pooled from two or three independent experiments. Data in **A, B, and G** are presented as mean ± s.e.m. Data in **E, F, I, and J** are presented as mean ± s.d. The statistics were obtained by one-way ANOVA coupled with Dunnett’s multiple-comparisons test **(E, F, I, and J)**.

To better mimic *in vivo* conditions, we next assessed the efficacy of these constructs in human spleen organoids stimulated with live-attenuated influenza vaccine (LAIV), an extension of a previously reported tonsil organoid system that recapitulates key features of GC reaction (e.g., light zone/dark zone organization, somatic hypermutation, affinity maturation, and antigen-specific antibody production)^42,43^ (**Fig. 3C**). TGM1-hCD4 and TGM1-hCD8 selectively and potently reduced the corresponding LAIV-induced CD38⁺Ki-67⁺ CD4⁺ or CD8⁺ T cell populations, as reflected by decreased frequencies within both the targeted T cell subsets and total CD3⁺ T cells, whereas the targeting arms alone and low-affinity TGM1 had no effect (**Fig. 3D–F, and S6H)**. In contrast, hTGFβ1 did not potently suppress these populations, likely due to a systemic “sink effect” arising from its broad and non-specific activity (**Fig. 3D–F**). Moreover, plasmablast (PB) frequencies among B cells and influenza (flu)–specific IgG levels were unchanged by either human T cell–targeted TGM1 construct at both day 4 and day 7 **(Fig. S6I and S6J)**. Together, these findings demonstrate that TGM1-hCD4 and TGM1-hCD8 specifically suppress antigen-stimulated human CD4⁺ and CD8⁺ T cell activation and proliferation, but are insufficient to modulate B cell responses in human spleen organoids.

### Human B-cell-targeted TGF-β agonist selectively silences antigen-stimulated B cell activation and antibody production in human spleen organoids

Given the insufficient B-cell silencing by the T cell-targeted TGF-β agonists in human spleen organoids, we developed a human B-cell-targeted TGF-β agonist by fusing the low-affinity TGM1 domains 1–3 to a high-affinity human CD19 scFv^44,45^ (TGM1-hCD19) **(S7A and S7B)**, which induced pSMAD2/3 reporter activity with an approximately five orders of magnitude higher affinity than untargeted TGM1 (**Fig. 3G and S7C)**. We then sorted naïve and memory B cells from PBMCs of healthy donors (n=4) and SLE patients (n=2) and cultured them with CD40L and IL-21 in the presence of the indicated proteins or hTGFβ1 (**Fig. S7D**). The CD19 scFv did not affect the intracellular staining level of CD19 during flow cytometry analysis (**Fig. S7E**). TGM1-hCD19 or hTGFβ1 significantly reduced the absolute number of CD3^-^CD19^+^ B cells and CD3^-^CD19^+^CD20^dim^CD38^high^ PBs at the end of culture compared with PBS- or untargeted TGM1-treated cells (**Fig. S7E-G**). Lastly, we assessed the effects of TGM1-hCD19 on LAIV-induced B-cell antibody responses in human spleen organoids (**Fig. 3C**). The intracellular CD19 staining levels were intact even in the presence of CD19 scFv (**Fig. S7H**). TGM1-hCD19 significantly reduced the percentage of CD3^-^CD19^+^ B cells in CD45+ leukocytes (**Fig. S7H**). TGM1-hCD19 also significantly suppressed LAIV-induced maturation of CD3^-^CD19^+^CD20^dim^CD38^high^ PBs (**Fig. 3H and 3I**). PBs in our gating scheme were also Ki67^+^CD27^bright^IRF4^+^BLIMP1^+^ (**Fig. S7I**). Notably, hTGFβ1 failed to suppress PB development in this system (**Fig. 3H and 3I**), likely due to pronounced “sink” effects from abundant non–B cell populations within organoids. Finally, TGM1-hCD19 significantly inhibited LAIV-induced influenza-specific IgG secretion compared with LAIV alone or combined with untargeted TGM1 or hCD19 scFv (**Fig. 3J**). In contrast, hTGFβ1 only modestly suppressed IgG secretion at day 4 and had no significant effect by day 7 (**Fig. 3J**). Consistent with a pronounced sink effect, re-administering hTGFβ1 at the day-4 medium change did not further reduce flu-specific IgG secretion at day 7 (**Fig. S7J**), indicating that the limited activity of the non-targeted ligand cannot be overcome by replenishing it. Collectively, these results demonstrate that TGM1-hCD19 silences antigen-driven B-cell maturation into PBs and antibody production in sorted human B cells and spleen organoids.

### Human B-cell-targeted TGF-β agonist blocks B-cell differentiation trajectory toward antibody-secreting cells

To gain deeper insights into TGM1-hCD19-mediated suppression of antibody responses, we performed single-cell RNA sequencing (scRNA-seq) on B cells in human spleen organoids of two donors (**Fig. 4A**). We performed unsupervised Louvain clustering on 163,632 cells surviving quality control (QC), with curated cell-type annotations transferred from a published human tonsil atlas^46^ (**Fig. 4B**). We focused on B-cell lineage cells (130,987 cells in total), identifying distinct clusters representing activated B cells (expressing *BCL2A1* and *MYC*), GC B cells (intermediate *CD38*, *BCL2A1*, and *MYC* with high *MKI67*), PBs (*MKI67*, *IGHM/A/G*, and *JCHAIN*) and plasma cells (PCs) (*IGHM/A/G* and *JCHAIN* without *MKI67*) (**Fig. 4C**). LAIV increased PBs by ∼3-fold and PCs by ∼10-fold compared with mock organoids, while TGM1-hCD19 nearly eliminated PB/PCs (**Fig. 4D and 4E**). Importantly, TGM1-hCD19 increased naïve B-cell frequency by 2∼3-fold, while reducing activated B cells and GC B cells (**Fig. 4E and S8A)**, suggesting that B-cell-targeted TGF-β signaling actually rewires the entire B-cell population toward a naïve-like state rather than simply reducing antibody-secreting cells (ASCs). Pseudobulk aggregation followed by PCA revealed that TGM1-hCD19 did not appreciably shift transcriptional programs of non-PB/PC B cells (i.e., naïve to GC B cells) compared with mock, LAIV alone, and LAIV plus untargeted TGM1 (**Fig. 4F**). Conversely, PB/PC transcriptional programs differed between mock and LAIV-stimulated organoids, with TGM1-hCD19 rendering PB/PCs more similar to mock organoids (**Fig. 4F**). Consistently, gene-regulatory network analysis^47^ showed that TGM1-hCD19 reduced IRF4 and BLIMP1 activity to baseline levels **(Fig. S8B)**. Finally, TGM1-hCD19 reduced *IGHG1/2/3/4* mRNA expression in PB/PCs to levels lower than mock, LAIV-, and LAIV plus untargeted TGM1-treated organoids **(Fig. S8C)**. Thus, TGM1-hCD19 imposes quantitative contraction of ASCs and represses IgG expression in the residual ASCs.

**Figure 4.**
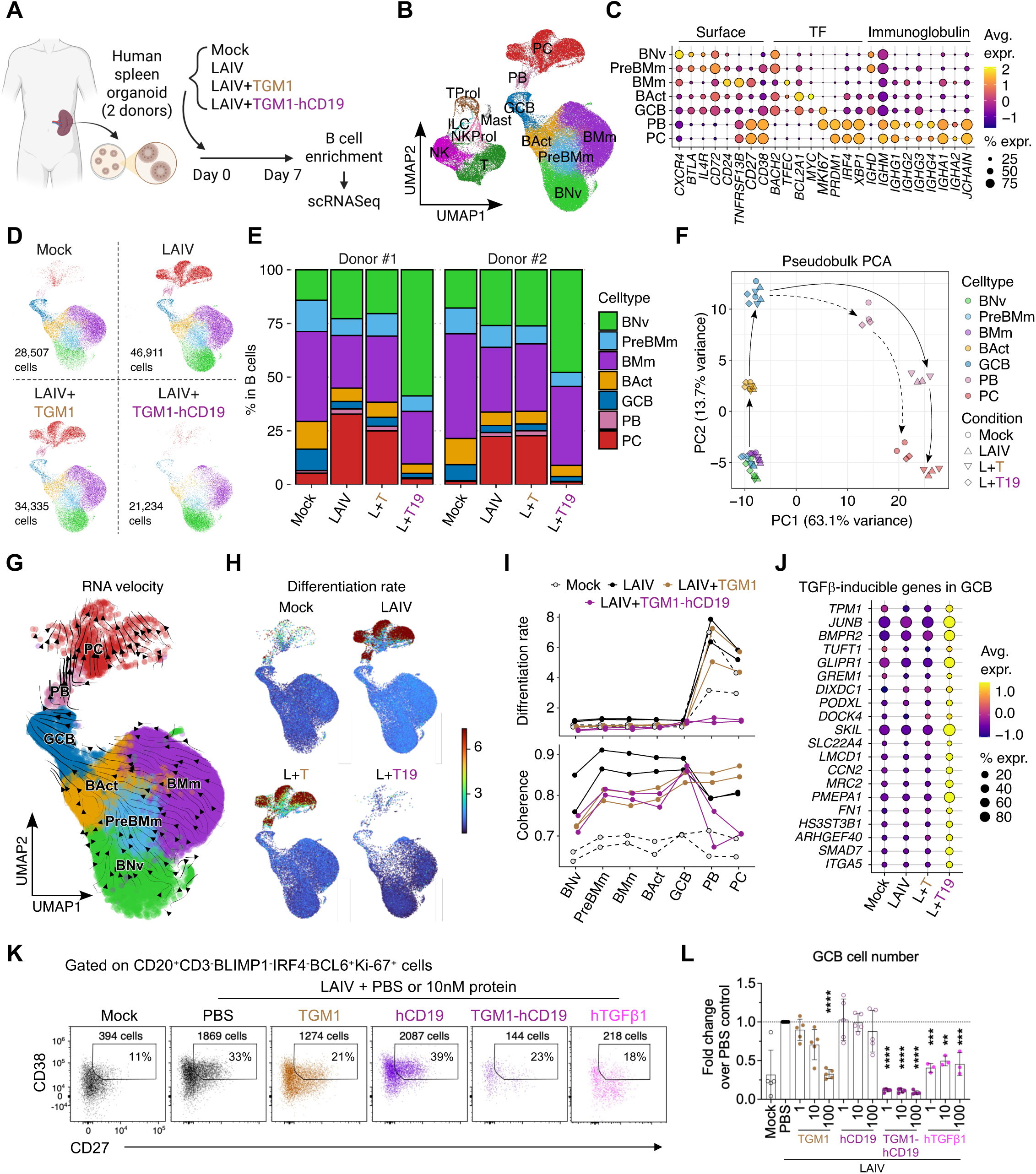
Human B cell-targeted TGF-β agonist suppresses antigen-driven B-cell differentiation in human spleen organoids. **A.** Schematic diagram illustrating the experimental design for scRNA-seq B cell sample preparation. **B.** UMAP visualization of B cell clusters from all samples combined, colored by annotated cell types. **C.** Bubble plot showing expression of B cell marker genes across distinct B cell clusters. **D.** UMAP visualization of B cell clusters across indicated treatment groups. **E.** Stacked bar plots showing the frequencies of distinct B cell subsets in indicated treatment groups across donors. **F.** PCA plots showing differentiation trajectories and transcriptional profiles of distinct B cell clusters across indicated treatment groups. **G.** RNA velocity stream plots of distinct B cell clusters in mock-treated samples. **H.** Estimated differentiation rates (cell state transitions) in indicated treatment groups. **I.** Differentiation rates and coherence of RNA velocity vectors across distinct B cell clusters in indicated treatment groups. **J.** Bubble plot showing expression of leading-edge genes of the TGF-β pathway enriched in GC B cells treated with LAIV plus CD19-targeted TGM1 across indicated treatment groups. **K.** Representative flow cytometry plots showing GC B cell frequencies and absolute counts in indicated treatment groups. **L.** Quantification of normalized GC B cell counts in the indicated treatment groups. Data in **K and L** are pooled from three independent experiments. Data in **L** are presented as mean ± s.d. The statistics were obtained by one-way ANOVA coupled with Dunnett’s multiple-comparisons test **(L)**.

To understand the critical point where TGF-β signaling blocks B-cell differentiation toward PB/PCs, we next performed RNA velocity analysis^48^. The RNA velocity vector field, embedded in the UMAP space (**Fig. 4G**), enabled inference of differentiation rates (i.e., the magnitude of dynamic change in the cell’s global mRNA transcription pattern) and coherence (i.e., concordance of the cell’s velocity vector with its neighbors) in each cell (**Fig. 4H and 4I**). The differentiation rates, which drastically increased at the PB stage in control organoids, remained nearly zero in TGM1-hCD19-treated organoids (**Fig. 4H and 4I**), indicating a profound silencing of GC B-to-PB differentiation. In the TGM1-hCD19-treated organoids, RNA velocity coherence remained high from naïve to GC B-cell stages but dropped at the PB stage (**Fig. 4I**), indicating a loss of differentiation “directionality” during the GC B-to-PB transition. To understand gene expression programs that govern the TGM1-hCD19-mediated silencing of GC B-to-PB transition, we performed several pathway-specific analyses. As expected, TGF-β signature scores were high across all B-cell subsets, including GC B cells, in the TGM1-hCD19-treated organoids (**Fig. 4J and S8D)**. Notably, GC B cells expressed the TGF-β signature genes at the lowest levels among B-cell subsets across conditions (**Fig. S8D**), suggesting a critical inhibitory role for TGF-β signaling at this stage. TGM1-hCD19 modestly increased senescence signature scores at later differentiation stages, without affecting apoptosis pathway genes **(Fig. S8D)**. We did not find substantial differences in cell-cycle states in any B-cell differentiation stages across conditions (**Fig. S8E and S8F**). Collectively, these observations suggest that B-cell-targeted TGF-β signaling imposes a qualitative phenotypic change at the GC B cell stage that abrogates their subsequent differentiation into ASCs.

To validate the impact of TGM1-hCD19 on GC B cells, we first re-analyzed the naïve B cell differentiation assay data (**Fig. S7D and S9A**). We found a striking contraction of GC B cells (CD27^+^Ki-67^+^BCL6^+^C-MYC^+^ non-PB B cells) in the presence of TGM1-hCD19 or hTGFβ1 (**Fig. S9B**), much more evident than the reduction of total B cell number (**Fig. S7F**). We also found that TGM1-hCD19 significantly reduced CD38 expression in the non-PB compartment (**Fig. S9C**), indicating a reduction in CD38^+^ GC B cells, in the spleen organoid LAIV stimulation assay. Indeed, TGM1-hCD19 significantly reduced the number of GC B cells (CD27^+^CD38^+^Ki-67^+^BCL6^+^ B cells after excluding BLIMP1^hi^IRF4^hi^ PBs) even at 1nM, while hCD19 did not cause any reduction (**Fig. 4K, 4L and S9D**). TGM1 at a high concentration (100 nM) and hTGFβ1 at all concentrations (1, 10, and 100nM) caused a ∼2-fold reduction of the GC B cell numbers (**Fig. 4L**), contrary to their minimal effects on the development of PBs (**Fig. 3I**), suggesting that GC B cells are more sensitive to TGF-β signals than PBs. Finally, we found that residual GC B cells in the presence of TGM1-hCD19 had lower BLIMP1 and C-MYC expression (**Fig. S9E**). BLIMP1 is lowly expressed in normal GC B cells and promotes the exit from GCs (i.e., GC B-to-PB transition^49^). The C-MYC protein governs the proliferative capacity of GC B cells^50^. Thus, our data show that B-cell-targeted TGF-β signaling suppresses the development of GC B cells and their maturation into ASCs.

### Therapeutic efficacy of mouse CD4⁺ T cell-targeted TGF-β agonist in CD4⁺ T cell–driven neuroautoimmune model

We next evaluated the therapeutic efficacy of T cell–targeted TGF-β agonists in mouse disease models. We first focused on a model of autoimmune neuroinflammation, experimental autoimmune encephalomyelitis (EAE) induced by MOG_35-55_. Although substantial studies have examined the role of TGF-β in EAE progression, its function remains controversial. Due to its dual capacity to suppress T cell activation and expansion while promoting Th17 differentiation, the net effect of TGF-β in EAE is context- and dose-dependent and can lead to either disease attenuation^51–55^ or exacerbation^22,56,57^. We immunized mice with MOG_35-55_ emulsified in CFA and administered the indicated proteins every other day starting on day 0 and continuing throughout the disease course (**Fig. 5A**). Our results showed that TGM1–mCD4 effectively prevented the development of EAE, with only 1 of 21 mice developing disease, whereas neither untargeted TGM1 nor TGM1–mCD8 provided significant therapeutic benefit, with EAE developing in ≥14/21 mice in the other groups (**Fig. 5B and 5C**). Flow cytometric analysis further showed that TGM1–mCD4 treatment substantially reduced CD4⁺ T cell numbers and the frequency of CD4⁺ T cells among total T cells in the spinal cord (SC) (**Fig. 5D, S10A, and S10B)**. Analysis of CD4⁺ T cell phenotypes in the SC revealed that TGM1-mCD4 did not significantly alter the frequencies of Foxp3⁺ or RORγt⁺ cells within the CD4⁺ T cell compartment, whereas their absolute numbers were substantially reduced **(Fig. S10C-E)**. In addition, the frequency of IFN-γ–producing CD4⁺ T cells was decreased, while the frequency of IL-17A–producing cells was largely unchanged **(Fig. S10F)**; however, the absolute numbers of both populations were markedly reduced (**Fig. 5E**). Notably, both the frequency and number of GM-CSF–producing CD4⁺ T cells in the SC were significantly reduced by TGM1-mCD4 (**Fig. 5F**), indicating effective suppression of pathogenic Th17-associated effector function. Together, these data demonstrate that TGM1-mCD4 effectively suppresses MOG-induced neuroinflammation, likely through potent and selective activation of TGF-β signaling that inhibits inflammatory CD4⁺ T cell expansion and accumulation rather than promoting pathogenic Th17 polarization.

**Figure 5.**
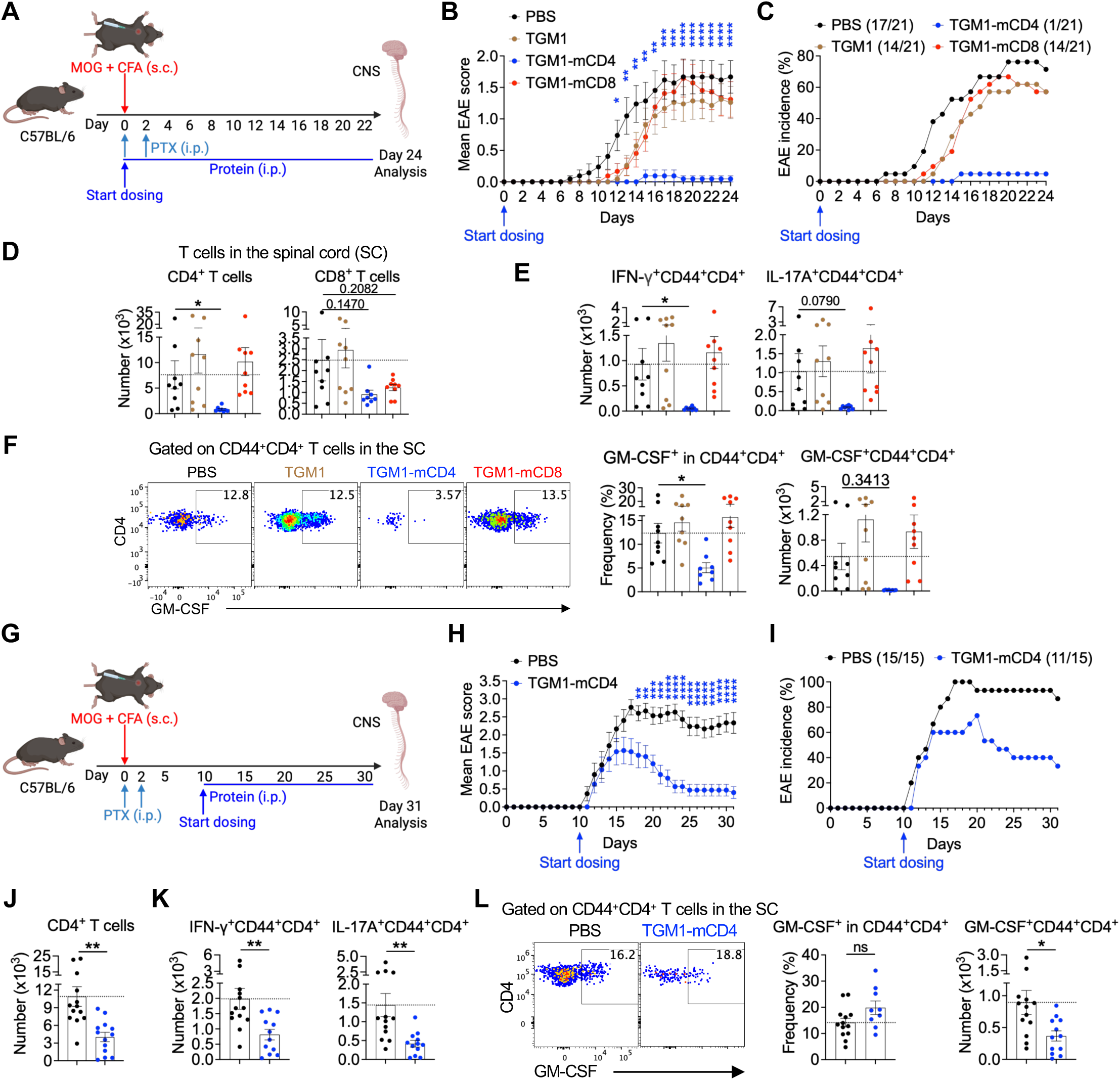
Therapeutic efficacy of mouse CD4⁺ T cell-targeted TGF-β agonist in the MOG₃₅–₅₅-induced EAE model. **A.** Schematic diagram illustrating the experimental design of MOG_35–55_ immunization, pertussis toxin (PTX) administration, and protein treatment starting on day 0. **B.** Mean EAE clinical scores of mice from the indicated groups (n = 21 mice per group). **C.** EAE incidence in mice from the indicated groups (n = 21 mice per group). **D.** Quantification of CD4⁺ and CD8⁺ T cell numbers in spinal cords (SCs) of mice from the indicated groups (n = 9 mice per group). **E.** Quantification of IFN-γ– and IL-17A–producing CD44⁺CD4⁺ T cell numbers in SCs of mice from the indicated groups. **F.** Representative flow cytometry plots and quantification of GM-CSF–producing CD4⁺ T cell frequencies and numbers in SCs of mice from the indicated groups. **G.** Schematic diagram illustrating the experimental design of MOG_35–55_ immunization, PTX administration, and protein treatment starting on around day 10 (n = 15 mice per group). **H.** Mean EAE clinical scores of mice in the indicated groups. **I.** EAE incidence in mice from the indicated groups. **J.** Quantification of CD4⁺ T cell numbers in SCs of mice from the indicated groups. **K.** Quantification of IFN-γ– and IL-17A–producing CD44⁺CD4⁺ T cell numbers in SCs of mice from the indicated groups. **L.** Representative flow cytometry plots and quantification of GM-CSF–producing CD4⁺ T cell frequencies and numbers in SCs of mice from the indicated groups. Data in **B-F and H-L** are representative of two independent experiments. Data are presented as mean ± s.e.m. The statistics were obtained by two-way ANOVA coupled with Šídák’s multiple comparisons test **(B and H)**, one-way ANOVA coupled with Dunnett’s multiple-comparisons test **(D-F)**, or unpaired Welch’s t-test (two-tailed) **(J-L)**.

In addition to the prophylactic setting, we next assessed the efficacy of TGM1-mCD4 in established disease by initiating protein treatment around day 10, when the first clinical signs of EAE typically begin to emerge and MOG-specific CD4⁺ T cells have already been activated (**Fig. 5G**). Under these conditions, TGM1-mCD4 treatment reduced peak disease severity, accelerated disease remission, and decreased disease incidence compared with the PBS control group (**Fig. 5H and 5I**), demonstrating that CD4⁺ T cell-targeted TGF-β signaling can exert therapeutic activity at or even after the onset of neuroinflammation in EAE. Flow cytometric analysis of the SC further revealed that TGM1-mCD4 treatment significantly reduced the accumulation of CD4⁺ T cells in the central nervous system (CNS) (**Fig. 5J**). Although the frequencies of IFN-γ⁺ and IL-17A⁺ cells among infiltrating CD4⁺ T cells were not significantly altered, their absolute numbers were markedly reduced following TGM1-mCD4 treatment (**Fig. 5K and S10G)**. Similarly, although the frequency of GM-CSF⁺ CD4⁺ T cells was not significantly decreased, their absolute number was significantly reduced (**Fig. 5L**). Together, these results indicate that, even in the context of ongoing disease, TGM1-mCD4 reduces disease severity and promotes disease remission, potentially through limiting the accumulation of pro-inflammatory CD4⁺ T cells in the CNS.

### Therapeutic efficacy of mouse CD4⁺ T cell-targeted TGF-β agonist in CD4⁺ T cell–driven airway inflammation model

Next, we further evaluated the therapeutic efficacy of T cell–targeted TGF-β agonists in OVA-induced type II airway inflammation. Mice were sensitized with OVA emulsified in aluminum hydroxide (alum) three times at weekly intervals, followed by three intranasal OVA challenges beginning one week after the final sensitization (**Fig. 6A**). The indicated proteins were administered intraperitoneally every other day starting on day 0 and continuing throughout the challenge phase (**Fig. 6A**). The data showed that TGM1-mCD4 effectively ameliorated lung inflammation, as evidenced by reduced histological inflammation scores and decreased eosinophil infiltration in both the bronchoalveolar lavage fluid (BALF) and lung compared with PBS controls (**Fig. 6B, 6C, S11A, and S11B)**. In addition, serum levels of OVA-specific IgG1, total IgE, and OVA-specific IgE were substantially reduced (**Fig. 6D and S11C)**. In contrast, low-affinity TGM1 and TGM1-mCD8 showed no evident therapeutic effect in this model (**Fig. 6B-D, S11B, and S11C)**. Analysis of lung-infiltrating T cells showed that TGM1-mCD4 potently reduced CD4⁺ T cell numbers and their frequency within total T cells in the lung (**Fig. 6E and S11D)**. Notably, TGM1-mCD4 decreased the proportion of Gata3⁺ cells while increasing Foxp3⁺ cell frequencies **(Fig. S11E and S11F)**, suggesting suppression of Th2 differentiation with a concomitant shift toward a regulatory phenotype. However, absolute numbers of both Gata3⁺ and Foxp3⁺ CD4⁺ T cells were reduced in the lung **(Fig. S11G)**, consistent with overall inhibition of CD4⁺ T cell expansion and infiltration. Importantly, IL-4 production by CD4⁺ T cells was significantly suppressed by TGM1-mCD4, as evidenced by reductions in both frequency and cell numbers in the lung (**Fig. 6F**). Together, these data demonstrate that CD4⁺ T cell–targeted TGM1 effectively attenuates type II airway inflammation likely by limiting CD4⁺ T cell expansion and accumulation and suppressing Th2 effector differentiation.

**Figure 6.**
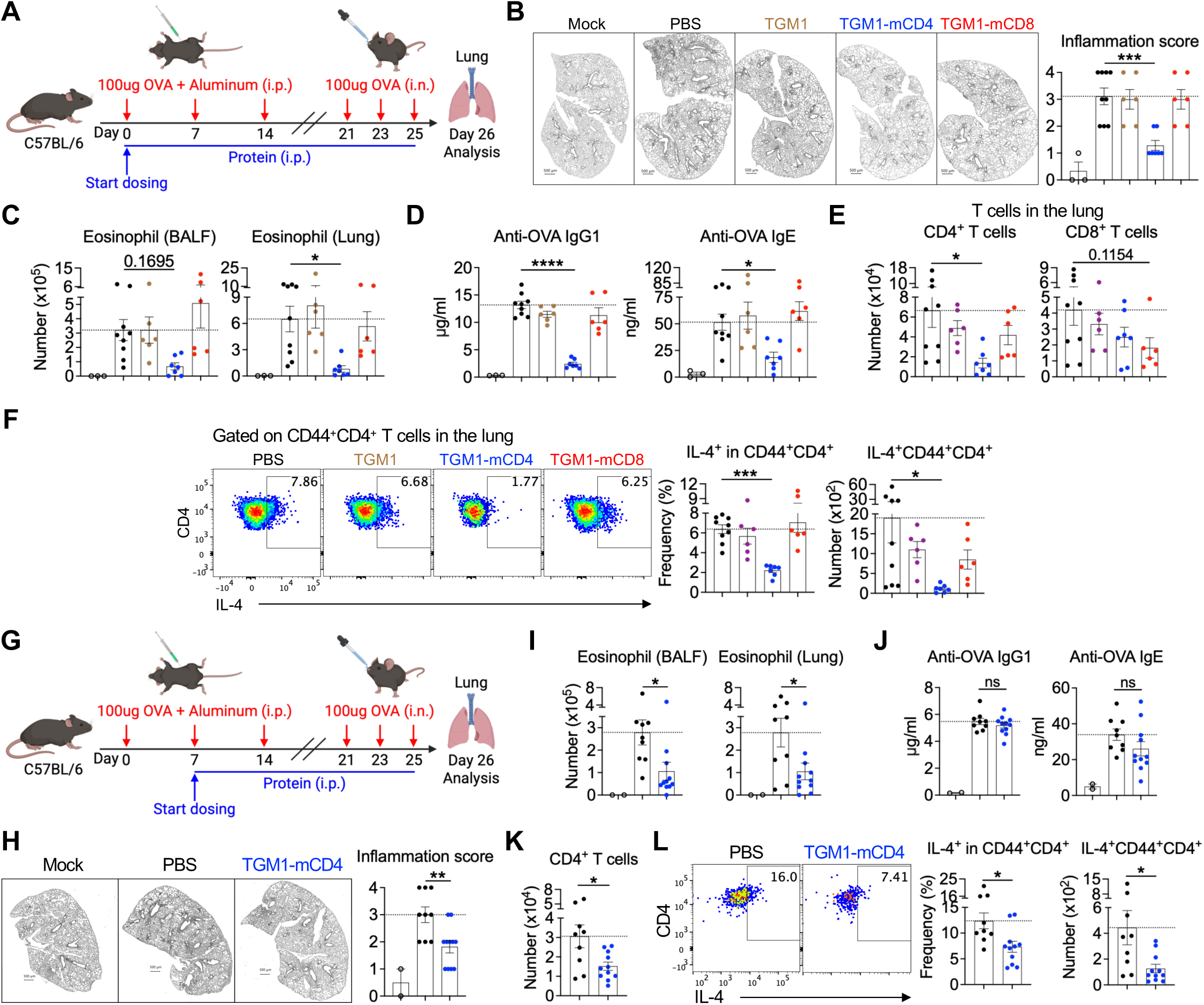
Therapeutic efficacy of mouse CD4⁺ T cell-targeted TGF-β agonist in the OVA-induced asthma model. **A.** Schematic diagram illustrating the experimental design of OVA challenge and protein administration starting on day 0 (Mock, n = 3 mice; PBS, n = 9 mice; TGM1, n = 6 mice; TGM1-mCD4, n = 7 mice; TGM1-mCD8, n = 6 mice per group). **B.** Representative lung histology and quantification of inflammation scores. Scale bars, 500 μm. **C.** Quantification of eosinophil numbers in BALF and lung tissue from mice of the indicated groups. **D.** Quantification of serum OVA-specific IgG1 and IgE levels in mice from the indicated groups. **E.** Quantification of CD4⁺ and CD8⁺ T cell numbers in lungs of mice from the indicated groups. **F.** Representative flow cytometry plots and quantification of IL-4–producing CD44^+^CD4⁺ T cell frequencies and numbers in lungs of mice from the indicated groups. **G.** Schematic diagram illustrating the experimental design of OVA challenge and protein administration starting on day 7 (Mock, n = 2 mice; PBS, n = 9 mice; TGM1-mCD4, n = 11 mice per group). **H.** Representative lung histology and quantification of inflammation scores. Scale bars, 500 μm. **I.** Quantification of eosinophil numbers in BALF and lung tissue from mice of the indicated groups. **J.** Quantification of serum OVA-specific IgG1 and IgE levels in mice from the indicated groups. **K.** Quantification of CD4⁺ T cell numbers in lungs of mice from the indicated groups. **L.** Representative flow cytometry plots and quantification of IL-4–producing CD44^+^CD4⁺ T cell frequencies and numbers in lungs of mice from the indicated groups. Data in **B-F and H-L** are representative of two independent experiments. Data are presented as mean ± s.e.m. The statistics were obtained by one-way ANOVA coupled with Dunnett’s multiple-comparisons test **(B-F)** or unpaired Welch’s t-test (two-tailed) **(H-L)**.

To further evaluate the therapeutic efficacy of TGM1-mCD4 in OVA-induced asthma after the initiation of antigen-specific CD4⁺ T cell responses, we initiated TGM1-mCD4 treatment on day 7, at the time of the second OVA/alum sensitization, when OVA-specific CD4⁺ T cells had already been primed by the initial sensitization (**Fig. 6G**). TGM1-mCD4 treatment significantly reduced lung inflammation compared with the PBS control, as evidenced by decreased histological inflammation scores and reduced eosinophil accumulation in both the BALF and lung (**Fig. 6H, 6I, and S11H)**. However, treatment initiated on day 7 did not significantly reduce serum OVA-specific IgG1, OVA-specific IgE, or total IgE levels compared with the PBS control (**Fig. 6J and S11I)**. Importantly, TGM1-mCD4 treatment substantially reduced the accumulation of CD4⁺ T cells in the lung (**Fig. 6K**). Moreover, both the frequency and absolute number of IL-4-producing CD4⁺ T cells were significantly decreased (**Fig. 6L**), indicating suppression of the inflammatory type 2 CD4⁺ T cell response. Together, these results demonstrate that TGM1-mCD4 retains therapeutic efficacy when administered after antigen-specific CD4⁺ T cell priming, suppressing CD4⁺ T cell accumulation and Th2 responses in the lung and potentially thereby limiting eosinophilic infiltration and lung inflammation.

### Therapeutic efficacy of T cell–targeted TGF-β agonists in mouse and human graft-versus-host disease models

TGF-β has previously been shown to attenuate GVHD severity and prolong survival in studies using non-targeted TGF-β activation or TGF-β neutralization^30,58^. Here, we further investigate whether selective delivery of TGF-β signaling to CD4⁺ or CD8⁺ T cells can achieve enhanced therapeutic efficacy in GVHD models. First, we evaluated the therapeutic efficacy of mouse T cell–targeted TGF-β agonists in an acute murine allogeneic GVHD model with major histocompatibility complex (MHC) mismatch by adoptively transferring CD45.1⁺ splenic T cells together with T cell–depleted bone marrow cells from C57BL/6 donor mice into lethally irradiated CD45.2⁺ BALB/c recipient mice (**Fig. 7A**). Both TGM1–mCD8 and TGM1–mCD4 showed robust therapeutic efficacy in suppressing GVHD progression, as evidenced by improved survival, delayed and reduced GVHD scores, and reversal of body weight loss (**Fig. 7B, 7C, and S12A)**. In contrast, mice treated with PBS or untargeted TGM1 developed severe GVHD, characterized by high mortality, elevated clinical scores, and pronounced weight loss (**Fig. 7B, 7C, and S12A)**. Histological analyses of colon tissues further demonstrated that both TGM1–mCD8 and TGM1–mCD4 effectively reduced colonic inflammation (**Fig. 7D**). Flow cytometric analysis revealed that TGM1–mCD8 substantially reduced CD8⁺ T cell numbers in both spleen and colon, whereas TGM1–mCD4 markedly reduced CD4⁺ T cells and also partially decreased CD8⁺ T cells (**Fig. 7E and S12B)**. Notably, colonic but not splenic CD8⁺ T cells were highly activated and differentiated into CD44^+^Granzyme B–producing cytotoxic cells, and this population was strongly suppressed by TGM1–mCD8 and more modestly reduced by TGM1–mCD4, as evidenced by reductions in both frequency and absolute numbers (**Fig. 7F and 7G**). In addition, these T cell–targeted TGM1 agonists essentially reduced IFN-γ– and TNF-α–producing inflammatory T cell populations in the colon **(Fig. S12C-E)**. Together, these data demonstrate that both CD8⁺- and CD4⁺ T cell–targeted TGM1 effectively suppress murine allogeneic GVHD, with the CD8⁺ T cell–targeted construct exhibiting the strongest efficacy.

**Figure 7.**
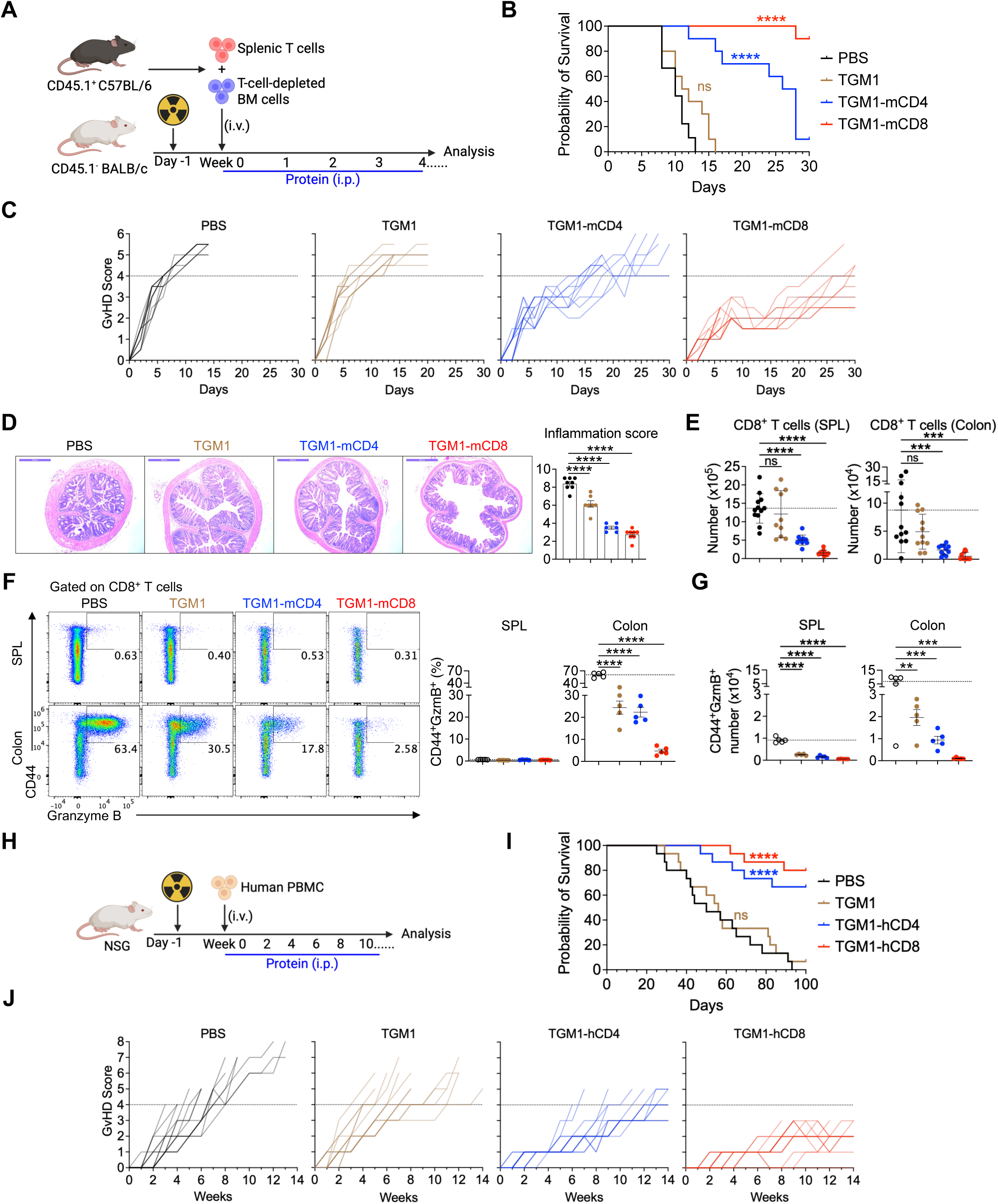
Therapeutic efficacy of T cell–targeted TGF-β agonists in mouse and human GVHD models. **A.** Experimental design of the mouse allogeneic GVHD model and protein administration (PBS, n = 14 mice; TGM1, n = 15 mice; TGM1-mCD4, n = 14 mice; TGM1-mCD8, n = 15 mice per group). **B.** Survival probability of mice from the indicated groups. **C.** Individual GVHD scores of mice from the indicated groups. **D.** Representative colon histology and quantification of colonic inflammation (PBS, n = 8 mice; TGM1, n = 7 mice; TGM1-mCD4, n = 6 mice; TGM1-mCD8, n = 8 mice per group). Scale bar, 1 mm. **E.** Quantification of CD8⁺ T cell numbers in spleen and colon of mice from the indicated groups. **F.** Representative flow cytometry plots and quantification of CD44⁺Granzyme B⁺ CD8⁺ T cell frequencies in the colon of mice from the indicated groups. **G.** Quantification of CD44⁺Granzyme B⁺ CD8⁺ T cell numbers in the colon of mice from the indicated groups. **H.** Experimental design of the human xenogeneic GVHD model and protein administration (n = 15 mice per group). **I.** Survival probability of mice from the indicated groups. **J.** Individual GVHD scores of mice from the indicated groups. Data in **B, C, E, I, and J** are pooled from two independent experiments. Data in **D, F, and G** are representative of two independent experiments. Data in **D, F, and G** are presented as mean ± s.e.m. Data in **E** are presented as mean ± s.d. The statistics were obtained by log-rank (Mantel–Cox) test **(B and I)** or one-way ANOVA coupled with Dunnett’s multiple-comparisons test **(D-G)**.

We further evaluated the efficacy of human T cell–targeted TGM1s in a humanized xenogeneic GVHD model by adoptively transferring human PBMCs into sublethally irradiated NSG recipients (**Fig. 7H**). Consistent with the murine allogeneic GVHD results, TGM1–hCD8 exhibited the strongest therapeutic efficacy, as evidenced by improved survival, reduced GVHD scores, and enhanced weight recovery, whereas TGM1–hCD4 conferred slightly reduced but still substantial benefit (**Fig. 7I, 7J, and S12F)**. In contrast, mice treated with PBS or untargeted TGM1 developed severe GVHD-related symptoms, characterized by high mortality, elevated clinical scores, and pronounced weight loss (**Fig. 7I, 7J, and S12F)**. Altogether, these data demonstrate that either CD8⁺- or CD4⁺ T cell–targeted TGF-β agonists provide significant therapeutic benefit in both murine allogeneic and humanized xenogeneic GVHD models.

## Discussion

Here, we demonstrate early-stage evidence that TGF-β agonism can achieve cell-specific silencing of antigen-driven adaptive T and B cell immune responses in mouse disease models and human spleen organoids. This approach does not induce cytotoxic cell death and is therefore unlikely to trigger adverse events such as CRS or ICANS. In addition, its high selectivity for targeted T or B cell subsets, with minimal effects on non-target populations, helps preserve the immune function of non-targeted immune populations, in contrast to broader immunosuppressive agents, although its specificity for truly pathogenic populations warrants further investigation. Furthermore, TGF-β signaling preferentially suppresses antigen-stimulated T and B cells during activation, thereby further maintaining overall immune homeostasis. Thus, this study suggests that a TGF-β agonism–based immune regulatory approach merits deeper investigation as a complement to existing immune cell–targeting therapeutics for the treatment of immunological diseases.

Previous studies of TGF-β function have largely relied on *in vivo* genetic models or *in vitro* systems. Our molecules provide alternative gain-of-function tools to dissect TGF-β signaling in defined immune subsets *in vivo* with high selectivity. For example, although previous *in vitro* studies and *in vivo* TGF-β receptor knockout models have shown that TGF-β is a key cytokine for Th17 differentiation and is essential for EAE induction^22,23^, we demonstrate a therapeutic and anti-inflammatory role of CD4⁺ T cell–specific delivery of strong TGF-β signaling using our CD4⁺ T cell–targeted TGM1. This results in effective suppression of pro-inflammatory and pathogenic CD4⁺ T cell differentiation and expansion, leading to superior therapeutic efficacy in preventing EAE onset, rather than increasing pro-inflammatory Th17 populations that drive disease. Moreover, this CD4⁺ T cell-targeted TGF-β agonist remained effective when administered after the establishment of inflammatory CD4⁺ T cell responses, reducing disease activity and promoting disease remission. These findings highlight its potential therapeutic efficacy in active Th17-driven disease.

Notably, previous studies have suggested that TGF-β plays a paradoxical role in either promoting or suppressing Tfh cell development, mainly based on *in vivo* TGF-β receptor knockout models or *in vitro* studies^59–63^. In this study, CD4⁺ T cell–specific delivery of strong TGF-β signaling demonstrated potent suppression of antigen-specific Tfh cell development and accumulation *in vivo*, resulting in a defective B cell response and antibody production. This inhibition of antigen-specific Tfh and humoral responses by TGM1-mCD4 may also contribute to reduced anti-drug antibody formation against TGM1 itself, as it is a helminth-derived non-self protein, thereby supporting its sustained efficacy *in vivo*. Therefore, TGF-β likely plays a complex, context- and dose-dependent role in regulating CD4⁺ T cell biology, where strong CD4⁺ T cell–intrinsic TGF-β signaling is primarily anti-inflammatory and holds significant therapeutic potential rather than being pro-inflammatory.

Additionally, the CD4⁺ T cell-targeted TGF-β agonist demonstrated protective efficacy in the mouse OVA-induced asthma model, suppressing Th2 responses and CD4⁺ T cell accumulation in the lung both when administered prophylactically and when treatment was initiated after antigen-specific CD4⁺ T cell priming. In both settings, these effects were accompanied by reduced eosinophilic infiltration and lung inflammation, consistent with the established role of Th2 cell responses in driving inflammatory cell recruitment to the lung^64^. However, treatment initiated after antigen sensitization did not significantly reduce serum IgG1 or IgE levels, in contrast to the prophylactic regimen, in which these antibody responses were substantially reduced. These findings suggest that, once an antigen-specific humoral response has been initiated, delayed targeting of CD4⁺ T cells may be less effective at suppressing the ongoing B cell response, potentially because B cell activation and differentiation have already progressed to a stage that is less dependent on continued CD4⁺ T cell help^65,66^. Given the multifaceted pathogenesis of asthma involving both pathogenic CD4⁺ T cell responses and B cell-mediated humoral immunity^67^, simultaneous targeting of CD4⁺ T cells and B cells with their respective cell-targeted immune silencers may provide a more comprehensive therapeutic strategy.

TGF-β critically contributes to the establishment of immune tolerance primarily through promoting Treg differentiation^68,69^. Our recent study showed that co-delivery of TGF-β and IL-2 signaling enables robust induction of antigen-specific pTregs *in vivo*, whereas TGF-β alone, in the absence of IL-2, fails to induce sufficient functional pTregs for immune tolerance^33^. Similarly, the present study shows that CD4⁺ T cell-targeted TGM1 alone does not induce a substantial Treg population in mouse models of OVA/CFA immunization, EAE, or asthma. This limited Treg induction may reflect the absence of a co-delivered IL-2 signal, the highly inflammatory environments generated by antigen immunization with CFA or alum, in which pro-inflammatory cytokines such as IL-6 and IL-1β antagonize TGF-β-driven pTreg differentiation while promoting effector T cell differentiation^70,71^, as well as the lack of highly tolerogenic conditions involving repeated high-dose intraperitoneal administration of soluble antigen in the absence of inflammatory adjuvants^33^. These suggest that its therapeutic effect is driven primarily by direct inhibition of effector programs and reprogramming into a non-inflammatory “quiescent” state, rather than robust Treg induction. Accordingly, this study examines a distinct strategy of direct effector lymphocyte silencing via cell type–selective TGF-β agonism without co-delivered IL-2, providing an alternative TGF-β–based therapeutic approach that is mechanistically distinct from IL-2–TGF-β combinatorial strategies. These two approaches are mechanistically orthogonal and may be complementary in different clinical contexts.

Moreover, we observed modest cross-suppression of CD4- and CD8-targeted TGF-β agonists on CD8⁺ and CD4⁺ T cells in mice, respectively. This effect may partially reflect the intrinsic low-affinity interaction of TGM1 with TGF-β receptors^33,34^, although indirect effects arising from functional crosstalk between CD4⁺ and CD8⁺ T cells could also contribute. CD4⁺ T cells provide important help for CD8⁺ T cell responses through multiple mechanisms, including the production of IL-2 and IL-21 and CD40L–CD40-mediated licensing of antigen-presenting cells^39,72,73^. Thus, suppression of OVA-specific CD4⁺ T cell activation by TGM1-mCD4 could potentially reduce these helper signals and thereby indirectly influence CD8⁺ T cell activation and expansion. Conversely, activated CD8⁺ T cells can also produce cytokines, including IL-2 and IL-21^74,75^, which may influence broader T cell responses. Such interdependent immune-cell responses could therefore contribute, at least in part, to the modest cross-suppression observed at high agonist concentrations. Additionally, TGM1-mCD8 did not significantly affect the abundance of CD8α⁺ DCs or their expression of CD80 and CD86 in OVA/CFA-immunized mice. Although the basis for the differential effects of TGM1-mCD8 on CD8⁺ T cells and CD8α⁺ DCs remains unclear, it may reflect cell type- and context-dependent responses to TGF-β signaling, which can vary according to DC subset, maturation state, and inflammatory environment^76^. These observations are consistent with, but do not establish, the possibility that the immunosuppressive effects of TGM1-mCD8 in this setting arise primarily from direct suppression of CD8⁺ T cells rather than substantial inhibition of CD8α⁺ DCs.

Anti-CD20 mAb-based B-cell depletion therapy has been in clinical use for over 20 years^10,11^, demonstrating tangible efficacy and a clinically tolerable side-effect profile. More recently, CD19 CAR-T cell therapy has shown superior efficacy in trials for refractory autoimmune diseases, including those resistant to anti-CD20 mAb therapy, owing to more profound B-cell depletion in deep tissues and effective targeting of CD19^+^CD20^dim^ plasmablasts^13,14,77^. Nevertheless, long-lasting plasma cells that lack CD19 expression persist and maintain preexisting humoral immunity to vaccine-related antigens even after CD19 CAR-T cell therapy^78^, suggesting that CD19 CAR-T cell therapy does not entirely erase the humoral immune memory. Limited long-term follow-up data are currently available for CD19 CAR-T-treated autoimmune patients. Given their substantially longer life expectancy compared to patients with malignancy, we anticipate that ancillary immunosuppressive therapeutics will remain necessary to manage disease relapses or exacerbations, even in the post-CD19-CAR-T era.

CD19-targeted TGF-β agonism, which acts primarily by silencing GC B cells poised to differentiate into PBs, represents an orthogonal approach to existing cytolytic-based B-cell-depleting therapeutics. Despite classical studies demonstrating the inhibitory effects of TGF-β on B cell proliferation and antibody secretion^80,81^, this pathway has received limited attention in contemporary molecular biology research, likely because TGF-β’s picomolar receptor affinity and pleiotropic effects have precluded its therapeutic application. Endogenous TGF-β is essential to promote GC B cell maturation from the light zone to the dark zone^82^. However, our data clearly show that exogenous TGF-β agonism inhibits B cell activity, most strikingly at the GC stage. Notably, the TGF-β signature gene expression was lowest at the GC B cell stage, suggesting that escape from TGF-β is necessary to maintain an appropriate GC B-cell differentiation program. Several non–mutually exclusive mechanisms could account for this. Activation-induced downregulation of TGF-β receptor expression, as demonstrated in TCR-stimulated T cells^79^, would render highly proliferative GC B cells transiently refractory to TGF-β; the BCL6- and c-MYC-driven GC program is functionally opposed to the growth-inhibitory and pro-differentiative actions of TGF-β and may actively restrain the pathway; and the availability of active ligand within the GC niche is itself constrained by integrin-dependent activation of latent TGF-β and by competition among the many receptor-bearing cells present. We favor the interpretation that this “TGF-β-low window” licenses onward differentiation toward ASCs, and that enforcing TGF-β signaling at this stage with TGM1-hCD19 closes the window, arresting cells in a GC-like state and preventing ASC formation, consistent with the selective collapse of the GC B cell-to-PB transition in our RNA velocity analysis and with the reduced BLIMP1 and c-MYC expression in residual GC B cells. Defining the precise molecular drivers of this downregulation, in particular distinguishing receptor regulation from transcriptional antagonism, is an important direction for future study. Theoretically, B-cell-targeted TGF-β agonism offers advantages over existing B-cell-depletion therapies, including off-the-shelf use without the harmful preconditioning required for CD19 CAR-T therapies and potential synergy with T-cell-targeted therapies (e.g., CD4^+^ T-cell-targeted TGF-β agonists). It is encouraging that B cells derived from SLE patients showed sensitivity to B-cell-targeted TGF-β equivalent to that observed in healthy controls. Future studies employing humanized mouse models and xenograft systems with human CD19-expressing cells will enable rigorous *in vivo* validation of therapeutic efficacy and mechanistic investigation in disease-relevant contexts.

Overall, this study provides proof-of-concept that targeted TGF-β agonists can precisely silence specific immune cell types with strong therapeutic potential in both mouse disease models and human immune organoids, highlighting cell-targeted TGF-β agonism as a versatile strategy for treating inflammatory, allergic, autoimmune, and transplant-related diseases.w

## Materials and methods

### Mice

Six-to ten-week-old female and male C57BL/6J (IMSR_JAX:000664), BALB/c (IMSR_JAX:000651), and NOD-scid IL2Rγnull (NSG) mice (IMSR_JAX:005557), as well as other strains as indicated, were purchased from The Jackson Laboratory. OT-II (IMSR_JAX:004194) and OT-I (IMSR_JAX:003831) mice were crossed with Thy1.1 mice (IMSR_JAX:000406) to generate OT-II × Thy1.1 and OT-I × Thy1.1 mice. All animals were housed in AAALAC-accredited facilities.

### Protein production

Recombinant proteins were cloned into the pD649 mammalian expression vector (ATUM), which encodes an HA secretion signal peptide, an N-terminal MSA fusion, and a C-terminal 6×His tag. Expression constructs were transfected into Expi293F™ cells using the Expi293™ Expression System (Thermo Fisher Scientific). After 3–4 days of culture, proteins were purified from culture supernatants by Ni-NTA agarose (Qiagen) and further purified by size-exclusion chromatography on a Superdex™ 200 column using an ÄKTA system (Cytiva). Endotoxin was removed using the Proteus NoEndo HC Spin Column Kit (VivaProducts) and verified to be at acceptable levels with the Pierce™ Chromogenic Endotoxin Quant Kit (Thermo Fisher Scientific). Final preparations were formulated and concentrated in sterile PBS, flash-frozen in liquid nitrogen, and stored at −80 °C until use.

### Flow cytometry

The following antibodies were purchased from BioLegend: mouse CD45.2 (109839), CD3 (100206), CD4 (100453, 100430, 100428, 116004), CD8α (100706, 100804, 155022), CD8β (126616), TCRvα2 (127806, 127822), Thy1.1 (202528, 202522), CD25 (102012), CD44 (103026), CD69 (104543), CD11b (101259), GITR (126316), IL-17A (506928), IFN-γ (505832), Ki-67 (151212), TNF-α (506346), and IRF4 (646416); human CD3 (317324), CD4 (980806, 317408, 357410, 300526), CD8α (980904, 388808, 388704, 301049), CD25 (356134), FOXP3 (320126), CD86 (305432), IgM (314518), CD138 (356520), CD11c (337238), CD80 (305236), Ki-67 (350540), CD45 (304026), BCMA (357530), CD81 (349510), CD38 (303552), CD20 (302322), and HLA-DR (307674). The following antibodies were purchased from BD Biosciences: mouse Bcl6 (562401) and Rorγt (564722); human IgG (564229), CD21 (750614), Fas (752346), BAFFR (749942), CD10 (741825), CD23 (749448), IgD (562518), CD3 (560365), CD5 (566122), CD27 (566449), BCL6 (569142), CD19 (555413), and BLIMP1 (565274). The following antibodies and reagents were purchased from Invitrogen: mouse PD-1(48-9985-82), Foxp3 (12-5773-82, 17-5773-82), T-bet (25-5825-82), Gata3 (46-9966-42), IL-4 (17-7-41-82), and Fixable Viability Dye (65-0865-18). The following antibodies were purchased from Miltenyi Biotec: human IgA (130-113-480) and IRF4 (130-100-909). Ghost Red 780 Fixable viability dye was purchased from Cytek Biosciences (13-0865). Human C-MYC mAb was purchased from Santa Cruz Biosciences (sc-42 AF647).

For cell-surface staining, live cells were incubated with antibodies and viability dye in PBS at 4°C for 1 h, and dead cells were excluded based on viability dye staining. For transcription factor and cytokine staining, cells were fixed and permeabilized using the Foxp3/Transcription Factor Staining Buffer Set (00-5521-00, Invitrogen) and then stained intracellularly at room temperature for 2 h. For cytokine detection, cells were stimulated with Cell Stimulation Cocktail (00-4970-03, Invitrogen) and Protein Transport Inhibitor Cocktail (00-4980-03, Invitrogen) at 37°C for 5 h prior to fixation. Cells were resuspended in PBS and analyzed on a CytoFLEX flow cytometer (Beckman Coulter) or an Aurora cytometer (Cytek Biosciences). Data were analyzed using FlowJo v10.10.0.

For CD4 and CD8 staining following protein incubation, isolated primary mouse splenic CD4⁺ and CD8⁺ T cells, or recovered human CD4⁺ and CD8⁺ T cells from frozen PBMCs, were incubated with the indicated proteins in 100 μl PBS at 37 °C for 1 h. Cells were then washed twice with PBS and subsequently stained with flow cytometry antibodies at room temperature for 30 min.

### Phosphorylated SMAD Detection Assay

HEK-Blue™ TGF-β cells (hkb-tgfbv2, InvivoGen) were used for detection and quantification of TGF-β activity. Cells were cultured in DMEM (11960-044, Gibco) supplemented with 10% FBS (A5256-701, Gibco), 1% penicillin–streptomycin (15140-122, Gibco), and 100 µg/mL Zeocin (ant-zn-05, InvivoGen). Cells were plated in flat-bottom 96-well plates at 5 × 10⁴ cells per well in 200 µL medium and incubated overnight at 37°C to allow attachment. On the following day, the medium was replaced with fresh medium containing the indicated proteins or recombinant human TGFβ1 (781804, BioLegend) at the specified concentrations (final volume: 200 µL per well). After 24 h of incubation, 20 µL of supernatant was transferred to a new 96-well plate containing 180 µL of 1× QUANTI-Blue™ Solution (rep-qbs, InvivoGen) and incubated at room temperature for ∼3 h. Absorbance was measured at 620 nm using a plate reader (SpectraMax i3x).

Mouse NIH/3T3 fibroblasts (CRL-1658™, ATCC) and bEnd.3 endothelial cells (CRL-2299™, ATCC) were purchased from ATCC. For pSMAD2/3 staining, cells were serum-starved overnight in DMEM supplemented with 1% FBS, 1× GlutaMAX™ (35050-061, Gibco), 1% sodium pyruvate (11360-070, Gibco), and 1% penicillin–streptomycin. Cells were then stimulated with the indicated proteins or recombinant human TGF-β1 at the indicated concentrations for 25 min. Following stimulation, cells were fixed with Cytofix Fixation Buffer (554655, BD Biosciences), permeabilized with Phosflow™ Perm Buffer III (558050, BD Biosciences), and stained with an antibody against phosphorylated SMAD2 (pS465/pS467)/SMAD3 (pS423/pS425) (562696, BD Biosciences). pSMAD2/3 levels were quantified by flow cytometry.

### Mouse and human T cell isolation and *in vitro* culture

Mouse spleens and lymph nodes were harvested and mechanically dissociated to generate single-cell suspensions. Red blood cells were lysed with ACK lysis buffer (A10492-01, Gibco). CD4⁺ or CD8⁺ T cells were enriched by negative selection using the EasySep™ Mouse CD4⁺ T Cell Isolation Kit (19852, STEMCELL Technologies) or Mouse CD8⁺ T Cell Isolation Kit (19853, STEMCELL Technologies), respectively. Naïve T cells (CD44⁻CD25⁻) were then sorted on a Sony SH800S cell sorter; sorted populations consistently exceeded 99% purity. Human CD4⁺ or CD8⁺ T cells were isolated from frozen PBMCs using the EasySep™ Human CD4⁺ T Cell Isolation Kit (17952, STEMCELL Technologies) or Human CD8⁺ T Cell Isolation Kit (17953, STEMCELL Technologies). Mouse and human T cells were cultured in RPMI 1640 with GlutaMAX™ supplemented with 10% FBS, 10 mM HEPES (15630080, Gibco), 1% sodium pyruvate, 1% penicillin–streptomycin, and 0.1% 2-mercaptoethanol (21985-023, Gibco).

For mouse *in vitro* cultures, purified naïve T cells were plated at 0.75 × 10^6^ cells/mL in flat-bottom 96-well plates coated overnight with 5 μg/mL InVivoMAb anti-mouse CD3 (BE0002, Bio X Cell) and 2.5 μg/mL InVivoMAb anti-mouse CD28 (BE0015, Bio X Cell). Recombinant proteins were added at the indicated concentrations at culture initiation, and medium was refreshed with proteins on day 3. Cells were cultured at 37°C for a total of 5 days.

For human *in vitro* cultures, T cells were plated at 0.75 × 10^6^ cells/mL in flat-bottom 12-well plates coated overnight with 5 μg/mL InVivoMAb anti-human CD3 (BE0001, Bio X Cell) and 2.5 μg/mL InVivoMAb anti-human CD28 (BE0248, Bio X Cell) in the presence of 10 ng/mL recombinant human IL-2 (202-IL, R&D Systems) for 3 days. Cells were then re-plated at 0.75 × 10^6^ cells/mL in flat-bottom 96-well plates coated overnight with anti-human CD3 (5 μg/mL) and anti-human CD28 (2.5 μg/mL) and cultured with the indicated recombinant proteins. Medium was refreshed with proteins on day 3, and cells were cultured at 37°C for a total of 5 days.

### Mouse OVA immunization model

One million sorted Thy1.1⁺ naïve OT-II and OT-I cells were mixed and adoptively transferred into Thy1.1⁻ C57BL/6 recipient mice via retro-orbital intravenous injection. Starting 1-day post-transfer, mice were immunized with 100 μg OVA (A5503, Sigma-Aldrich) emulsified in complete Freund’s adjuvant (F5881-6X10ML, Sigma-Aldrich) and injected near the inguinal lymph nodes (ILNs). The indicated proteins were administered intraperitoneally at the doses specified in the corresponding figure legends every other day, starting on day 0. Mice were euthanized on day 7 or day 8 post-immunization, and inguinal lymph nodes and spleens were collected for analysis. For OT-II and OT-I bulk RNA-seq, T cells were enriched from inguinal lymph nodes on day 8, donor OT-II and OT-I cells were subsequently sorted, and samples were submitted to MedGenome, Inc. for library preparation and sequencing.

### Human B-cell sorting and *in vitro* stimulation

Cryopreserved PBMCs from four healthy donors and two patients with SLE were purchased from STEMCELL, Inc. Cryopreserved PBMCs from healthy donors were surface-stained in FACS buffer (PBS supplemented with 0.5% BSA and 2 mM EDTA), and naïve B cells (CD3^-^CD20^+^CD27^-^) were sorted by FACS. Cells were rested in complete medium overnight. The next day, cells were stimulated with MEGACD40L (Enzo, ALX-522–110-C010, 200 ng/mL) plus IL-21 (PeproTech, 200–21, 50 ng/mL) for 24 hours^80^. The next day, in-house proteins or recombinant human TGFβ1 (BioLegend) were added, and the cells were cultured for an additional 7 days. At the end of the culture, cells were harvested and analyzed by flow cytometry.

### Human spleen dissociation

Fresh spleen samples were obtained from deceased adult organ donors through Donor Network West (Northern California and Northern Nevada). The project was approved by the Donor Network West Ethics & Mission Committee. Spleens were decontaminated in Ham’s F-12 medium supplemented with Normocin and penicillin–streptomycin for at least 30 min at 4 °C. Tissues were transferred to Petri dishes, trimmed to remove large vessels, fat, and capsule, sectioned into ∼5 g pieces, and mechanically dissociated into a paste. Each section was resuspended in 10 mL dissociation medium consisting of complete RPMI (RPMI 1640 with GlutaMAX, 10% FBS, 1× penicillin–streptomycin, 1× non-essential amino acids, 1× sodium pyruvate, 1× insulin–transferrin–selenium, and 1× Normocin) supplemented with 1 mg/mL collagenase IV and 200 U/mL DNase I. The slurry was transferred to gentleMACS C-tubes and processed using two rounds of dissociation on a gentleMACS Dissociator (protocol Spleen_04_01). Cell suspensions were filtered through a 100 μm strainer, resuspended in PBS containing 2% FBS and 6 mM EDTA, and washed once. Red blood cells were lysed with ACK lysis buffer for 5 min at room temperature, and cells were washed, counted, and aliquoted into 50 mL conical tubes. Granulocytes and residual erythrocytes were depleted using the EasySep Direct Human PBMC Isolation Kit (STEMCELL Technologies) according to the manufacturer’s instructions. Cells were then counted, resuspended in freezing medium (FBS with 10% DMSO), aliquoted into cryovials, frozen at −80 °C, and transferred to liquid nitrogen (−150 °C) for long-term storage.

### Human spleen organoid culture and stimulation

Twelve-well plates were set up with one 12-mm Transwell insert (0.4-μm pore size) per well, and 1 mL organoid medium [complete RPMI with 0.5 μg/mL B-cell activating factor (BAFF)] was added to the lower chamber. Cryopreserved splenocytes were thawed and washed once in complete RPMI, counted, and resuspended at 6 × 10^7^ cells/mL. Aliquots of 100 μL splenocytes were transferred to 1.5-mL tubes, mixed with antigenic stimulus and in-house proteins, and dispensed into the Transwell inserts. For LAIV stimulation, 1 μL FluMist Quadrivalent (intranasal; 2022–2023 formulation) was added per well; mock controls received 1 μL PBS. In-house proteins were used at a final concentration of 10 nM. On day 3, 500 μL of the medium in the lower chamber was removed, and 600 μL of fresh organoid medium (without replenishing LAIV or in-house proteins) was added into the lower chamber. “Boost” hTGF-β1 indicates re-administration of rhTGF-β1 at the same concentration during the day 4 medium change. Organoids were harvested on day 7 for flow cytometry and single-cell RNA sequencing analyses. Supernatants were stored at −20 °C for ELISA.

### Human single-cell RNA-seq sample preparation

On day 7 of culture, human spleen organoids were dissociated by gently rinsing the Transwell membrane with FACS buffer (PBS supplemented with 0.5% BSA and 2 mM EDTA). B cells were enriched by negative selection using the Human Pan B Cell Isolation Kit (Miltenyi, 130-101-638) according to the manufacturer’s instructions. Cells were washed and resuspended in PBS containing 0.04% BSA. Single-cell capture was performed on a 10x Genomics Chromium X instrument, and gene expression libraries were generated using Single Cell 5′ R2-only v3 chemistry. Libraries were sequenced as paired-end 150-bp reads on an Illumina NovaSeq 6000. Reads were aligned to the GRCh38-2024-A reference transcriptome (including introns) and gene expression was quantified using Cell Ranger v9.0.1. Approximately 14,000–27,000 cells were recovered per sample.

### Mouse MOG_35–55_-induced EAE model

Female mice were immunized subcutaneously with 100 µg MOG₃₅–₅₅ peptide per mouse in complete Freund’s adjuvant (BD Biosciences) containing 200 µg Mycobacterium tuberculosis per mouse (BD Biosciences), injected at the axilla of both sides. Concurrently, 400 ng pertussis toxin per mouse (PTX, 181, List Labs) was administered intraperitoneally, with a second dose given 48 hours later. The indicated proteins (40–50 pmol per dose) were administered intraperitoneally every other day, starting on day 0 or around day 10, as specified in the corresponding figure legends. Mouse body weight and clinical signs of disease were recorded daily and scored according to the following scale: 1 – tail paralyzed; 2 – hind limb weakness; 3 – one or both hind limbs paralyzed; 4 – front limb paralysis; 5 – moribund or deceased.

### Mouse OVA-induced airway inflammation model

Mice received three weekly intraperitoneal injections of 100 μg OVA protein formulated in 150 μL Alhydrogel adjuvant (2%; vac-alu-50, InvivoGen). One week after the final adjuvant injection, mice were administered three intranasal doses of 100 μg OVA protein every other day. The indicated proteins (40–50 pmol per dose) were administered intraperitoneally every other day, starting on day 0 or day 7, as specified in the corresponding figure legends. Tissues were collected and analyzed one day after the final intranasal dose.

### Mouse allogeneic GVHD model

BALB/c female recipient mice received lethal total body irradiation (9 Gy). One day later, 5 × 10^6^ T cell–depleted bone marrow cells and 1 × 10^6^ splenic T cells isolated from donor female C57BL/6 mice were intravenously transferred into the BALB/c recipients by tail injection. The indicated proteins (40 pmol) were administered by intraperitoneal injection every other day starting on the day of cell transfer. Mice were monitored daily for clinical signs of GVHD, including weight loss, posture, mobility, and skin lesions. GVHD severity was scored according to standardized criteria **(Supplementary Table 1)**. Mice were humanely euthanized if the GVHD score exceeded 6 or if body weight loss reached ≥30% relative to baseline. Flow cytometry and histological analyses were performed approximately 10 days after cell transfer.

### Humanized xenogeneic GVHD model

10-to 12-week-old NSG mice received 2.4 Gy of whole-body X-ray irradiation and, 4–6 hours later, were intravenously injected with 3 × 10^6^ human PBMCs in 50 µL PBS, obtained from de-identified adult healthy blood donors. The indicated proteins (40 pmol) were administered by intraperitoneal injection every other day starting on the day of PBMC transfer. Mice were monitored daily for clinical signs of GVHD, including weight loss, lethargy, reduced mobility, hunching, ruffled fur, diarrhea, and skin lesions. GVHD severity was scored according to standardized criteria **(Supplementary Table 1)**. Mice were humanely euthanized if the GVHD score exceeded 8 or if body weight loss reached ≥30% relative to baseline.

### Isolation of immune cells from tissues

Mice were euthanized at the end of the indicated treatments. For spinal cord lymphocyte isolation, mice were perfused, spinal cords were collected, mechanically dissociated using the plunger of a 1 mL syringe and filtered through 70 μm strainers to obtain single-cell suspensions.

For BALF collection, lungs were flushed three times with 0.75 mL PBS via a catheter inserted into the trachea. For lymphocyte isolation from the lung, tissues were mechanically dissociated using the plunger of a 1 mL syringe and passed through 70 μm cell strainers to obtain single-cell suspensions.

For lamina propria lymphocyte isolation, colons were opened longitudinally, cut into ∼2 cm pieces, and incubated in 5 mM EDTA (AM9260G, Invitrogen) containing 1 mM DTT (R0861, Thermo Fisher Scientific) at 37 °C for 30 min to remove epithelial cells. Tissues were then minced and digested with DNase I (4 mg/mL, Roche) and collagenase D (0.5 mg/mL, Roche) at 37 °C for 30 min with shaking to generate single-cell suspensions, which were filtered through 70 μm strainers.

Lymphocytes were then enriched using a 40%/70% Percoll gradient (17089101, Cytiva), and cells at the interface were collected for flow cytometry analysis.

### ELISA

For human ELISA to detect anti-influenza antibodies, Costar plates were coated with 0.1 μg per well of the 2022–2023 Fluzone Quadrivalent inactivated influenza vaccine (Sanofi) and incubated overnight at 4 °C. Plates were washed with PBS containing 0.05% Tween-20 (PBST) and blocked with PBS containing 1% BSA for 2 h at room temperature. Plates were then washed and incubated with spleen organoid supernatants for 1 h at room temperature, washed again, and incubated with horseradish peroxidase–conjugated anti-human IgG Fc secondary antibody (SouthernBiotech) for 1 h at room temperature. Signal was developed with 1-Step Ultra TMB substrate (Thermo Fisher Scientific, 37574), stopped with ELISA Stop Solution (Thermo Fisher Scientific, SS04), and absorbance was measured at 450 nm using a 96-well plate reader (Cytation 7, Agilent).

For mouse ELISAs, blood samples were centrifuged at 3,000 × g and serum was collected. Serum antibody levels were quantified using commercial ELISA kits, including total IgE (ELISA MAX™ Standard Set Mouse IgE; BioLegend, 432401), OVA-specific IgE (LEGEND MAX™ Mouse OVA-Specific IgE ELISA Kit; BioLegend, 439807), and OVA-specific IgG1 (Mouse Anti-Ovalbumin IgG1 ELISA Kit; Cayman Chemical, 500830), according to the manufacturers’ instructions.

### Histology

Lung tissues from perfused mice and colon tissues were fixed in 10% neutral buffered formalin (Sigma-Aldrich, HT501128) and submitted to S. Avolicino for paraffin embedding, sectioning, and haematoxylin and eosin staining. Lung sections were imaged using a Keyence BZ-X810 microscope, whereas colon sections were imaged using a Leica DM2000 microscope. Histopathological evaluation was performed in a blinded manner.

Lung inflammation was scored on the basis of the extent of peribronchial and perivascular cellular infiltration: 0, no infiltrates; 1, a few inflammatory cells; 2, a one-cell-thick ring of inflammatory cells; 3, a 2–3-cell-thick ring; 4, a 4–5-cell-thick ring; 5, a ring >5 cells thick.

Colon inflammation was scored as follows: 0, no evidence of inflammation; 1, low-level inflammation with scattered infiltrating mononuclear cells (1–2 foci); 2, moderate inflammation with multiple foci; 3, high-level inflammation with increased vascular density and marked wall thickening; 4, severe inflammation with transmural leukocyte infiltration and loss of goblet cells.

### Bulk RNA-seq analysis

Reads were aligned to the mouse reference genome (mm10) using Rsubread (v.2.18.0). Gene expression was quantified with featureCounts. For principal component analysis and heatmap visualization, variance stabilizing transformation (VST) was applied to the raw read counts. Differential expression (DE) analysis was performed with DESeq2^81^. Genes with FDR-adjusted P value < 0.05 and |log2FC| > 1 were considered DE unless otherwise noted. Gene Set Enrichment Analysis (GSEA) was performed against the Hallmark gene sets using log2FC-based gene ranking as input. For gene co-expression module analysis (also known as weighted gene co-expression network analysis; WGCNA), VST-transformed read counts were initially split into 45 gene modules, among which 14 modules were significantly enriched in TGM1-mCD4-treated CD4+ T cells or TGM1-mCD8-treated CD8+ T cells, when compared with other treatment conditions. Enrichment analysis was performed against TF perturbation genesets using the Enrichr webtool (https://maayanlab.cloud/Enrichr/). For celltype signature analysis, transcripts per million reads (TPMs) were calculated for all genes, and gene symbols were converted to corresponding human orthologs using the ENSEMBL repository. TPM values of sort-purified T-cell subsets were retrieved from the Monaco datasets available in the Human Protein Atlas (https://www.proteinatlas.org/humanproteome/single+cell/immune+cell/data#immune_cells_monaco). The TPM values were converted into log2 scale with the addition of the pseudocount of 1. We then applied the EPIC algorithm to the log-transformed TPM values to infer the composition of T-cell subsets in our bulk RNAseq data, using the Monaco dataset as a reference^82^. The predicted probabilities for each cell type were log-transformed and scaled, and used as a proxy of the proportion of adoptively transferred OT-I and OT-II T cells that have acquired the specific phenotype to construct radar charts.

### Single-cell RNA-seq analysis

The CellRanger-filtered gene expression matrix was first processed using Seurat (v.5.3.1). For each dataset, we excluded cells with fewer than 200 genes (to remove debris, empty droplets, and low-quality cells) and cells with higher than 9000 genes or higher than 100,000 reads mapped to RNA (to remove large cell aggregates). We also excluded cells in which >10% of transcripts were derived from mitochondrial RNA, leaving 163,632 cells in total (33,179, 51,696, 43,858, 34,899 cells in mock-, LAIV-, LAIV plus TGM1-, and LAIV plus TGM1-hCD19-treated groups, respectively).

We then performed a standard Seurat preprocessing workflow, including log-normalization, variable gene selection, scaling, PCA, and Uniform Manifold Approximation and Projection (UMAP) with 10 PCs using default parameters. To annotate cell types, we first performed unsupervised Louvain clustering on K-nearest-neighbor graphs (k = 20, resolution = 1). We applied the Seurat celltype-label transfer workflow using Human tonsil atlas data (ver2; https://zenodo.org/records/8373756) as the reference. The annotations stored in the “annotation_figure_1” slot were used for transfer. Transfer anchors were established in the 30-dimensional PCA space using genes in our data that were also present in the reference data. Seurat-defined clusters were manually annotated based on canonical marker gene expression and the aggregated prediction scores returned from the label transfer workflow.

For pseudobulk PCA, gene expression was aggregated by donor, condition, and cell type using the AggregateExpression function in Seurat. Transcription factor (TF) activity prediction was conducted on B-cell data (downsampled to 10,000 cells per condition) using the pySCENIC (v.0.12.1) Docker distribution with the default parameter settings. A total of 265 regulons representing unique TFs, including IRF4 and BLIMP1, were identified. Normalized regulon activity scores were aggregated across donors, conditions, and cell types. Differential expression (DE) analysis was performed using Seurat’s FindMarkers function. Gene set enrichment analysis (GSEA) was performed using log2FC ranking as input against Hallmark and MSigDB C2 genesets (7,561 genesets in total) with the fgsea package. Leading-edge genes are selected for visualization as dot plots. For cell cycle gene signature scores, Seurats’s CellCycleScoring function was used with built-in S- and G2M-phase genes. For gene signature scores, Seurat’s AddModuleScore function was used. The scores were aggregated across donors, conditions, and cell types, and scaled for visualization.

For RNA velocity analysis, pre-mature (unspliced) and mature (spliced) transcript abundances were quantified with velocyto using position-sorted BAM files generated by the CellRanger pipeline, with filtering for cell barcodes that passed the Seurat QC step described above^48^. Outputs from velocyto were further analyzed using scVelo to estimate RNA velocity by stochastically modeling the transcriptional dynamics of splicing kinetics in individual cells^83^. The velocity vector fields were projected onto the UMAP space and visualized as velocity streams. The length of the velocity vector indicates the rate of differentiation (i.e., cell-state transition). The coherence of the vector field indicates how the differentiation pattern of a given cell correlates with that of its neighboring cells.

### Statistical analysis

Statistical analyses were performed using GraphPad Prism 10 (GraphPad Software). Data are presented as mean ± s.e.m. or mean ± s.d., as specified in the corresponding figure legends. P < 0.05 was considered statistically significant. Significance is denoted as *P < 0.05, **P < 0.01, ***P < 0.001, and ****P < 0.0001. Detailed statistical methods are provided in the corresponding figure legends. The number of mice per experiment is represented by individual data points in each figure and specified in the corresponding figure legends.

## Ethics statement

All experimental procedures were approved by the Stanford University Institutional Animal Care and Use Committee (IACUC; protocol IDs 32279, 34708, and 10269) and were conducted in accordance with institutional guidelines. Blood for PBMC isolation from healthy donors was provided by the Stanford Blood Center, which also obtained ethical approval for the donors.

## Acknowledgments

We acknowledge the Stanford Animal Care and Facilities at Stanford and the Stanford Cell Sciences Imaging Facility (RRID SCR_017787) at Beckman Center for their technical assistance. Schematics and diagrams were created using BioRender. We thank Lei Chen for assistance with human spleen processing.

## Author contributions

K. Christopher Garcia conceived the study, supervised the experiments, and collaborated with the authors on the manuscript. Qinli Sun designed and conducted the mouse and human T cell-targeted TGF-β agonist experiments, and Masato Ogishi designed and conducted the human B cell-targeted TGF-β agonist experiments. Qinli Sun and Masato Ogishi analyzed the data, prepared the figures, and wrote the manuscript. Elsa Solà and Jie Zhang performed the human spleen organoid experiments, analyzed the ELISA data, and processed the scRNA-seq samples. Alison K. Barrett established the EAE model, recorded clinical scores, and performed mouse perfusions. Hua Jiang designed and performed the mouse GVHD experiments and assisted with mouse tissue processing. Hao Yan designed and performed the humanized GVHD experiments. Ahmad Salehi assisted with human spleen sample preparation. Honghui Liu assisted with BALF extraction. Mark M. Davis supervised the human spleen organoid experiments. Tobias V. Lanz supervised the EAE experiments. Robert S. Negrin supervised the GVHD experiments. Huiyun Lyu, Peng Xiao, and Qizhi Tang contributed to the experimental design. All authors contributed to manuscript preparation.

## Data availability

The raw and processed bulk RNA-seq and scRNA-seq data generated in this study will be made publicly available in the GEO database upon manuscript acceptance. The R code used for RNA-seq data analysis will also be shared publicly at that time. Any additional information required to reanalyze the data reported in this paper is available from the lead contact upon request.

## Supplementary Figure Legends

**Figure S1.**
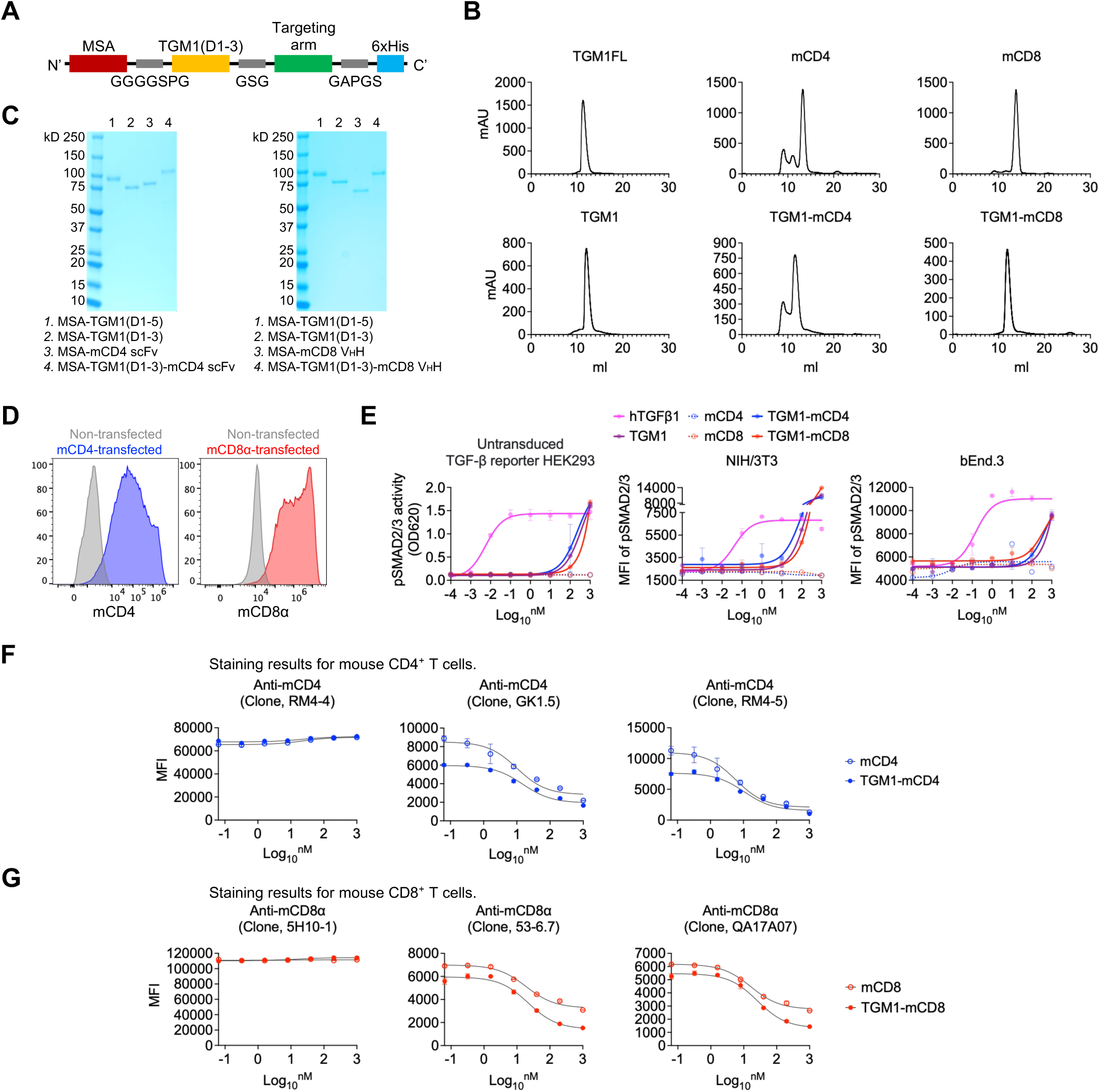
Design and characterization of mouse T cell-targeted TGF-β agonists. **A.** Schematic diagram of the plasmid construct encoding the MSA–TGM1–targeting arm fusion protein. **B.** Analytical size-exclusion chromatography (SEC) profiles of the indicated proteins after Ni-NTA affinity purification. The major elution peak represents the purified protein of interest. **C.** SDS–PAGE analysis showing the molecular weights and purity of the indicated recombinant proteins. **D.** Representative flow cytometry plots showing mouse CD4 and CD8α expression in mouse CD4- or CD8α-transduced TGF-β reporter HEK293 cells. **E.** Dose–response curves showing SMAD2/3 activation in untransduced TGF-β reporter HEK293 cells, measured using a pSMAD-responsive secreted alkaline phosphatase (SEAP) reporter assay, and in mouse NIH/3T3 fibroblasts and bEnd.3 endothelial cells, measured by pSMAD2/3 staining. **F.** Dose–response curves showing staining intensities of the corresponding commercial antibodies on primary CD4⁺ T cells after incubation with the indicated proteins. **G.** Dose–response curves showing staining intensities of the corresponding commercial antibodies on primary CD8⁺ T cells after incubation with the indicated proteins. Data in **B-G** are representative of two independent experiments. Data are presented as mean ± s.e.m.

**Figure S2.**
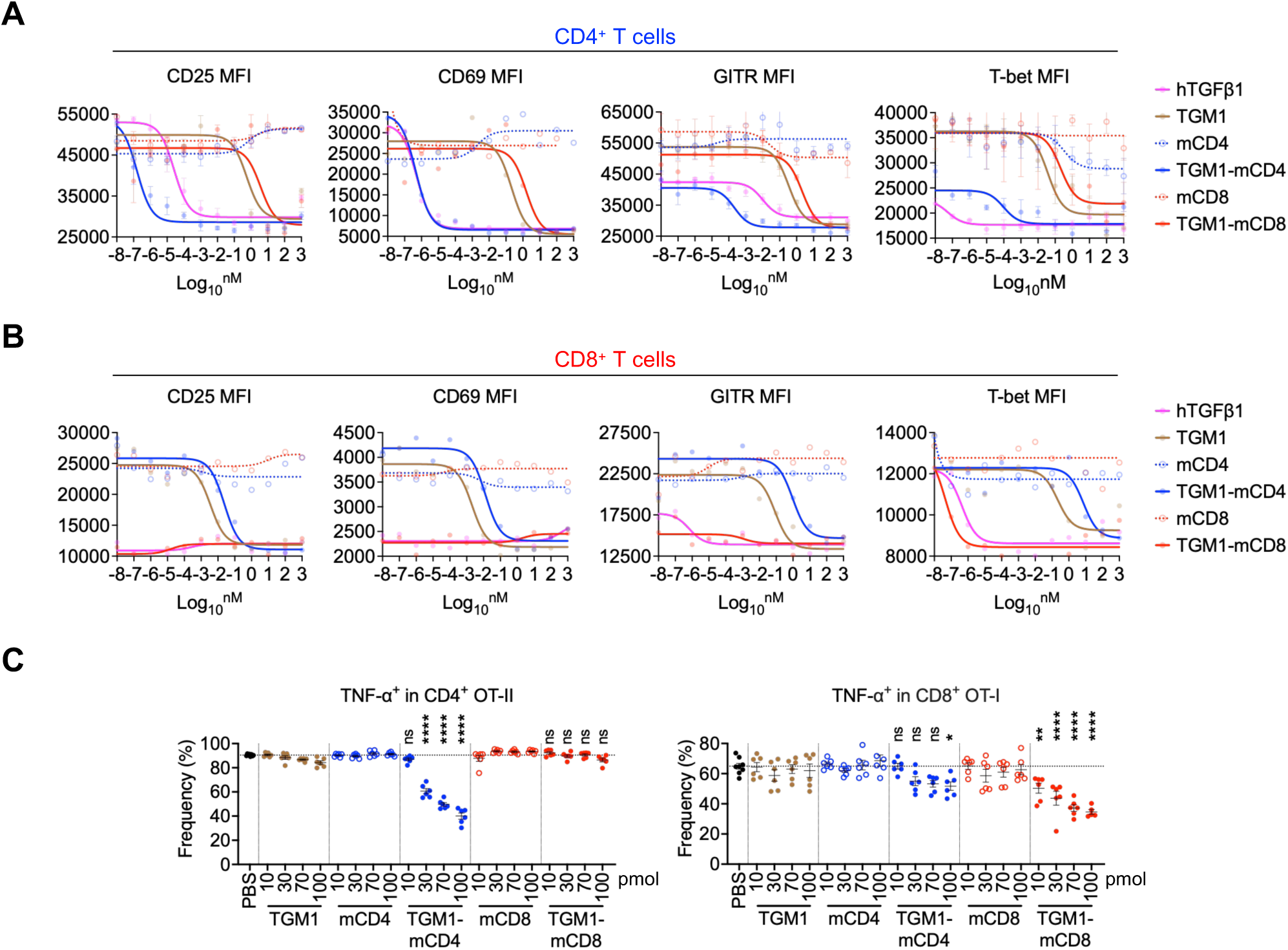
Effects of mouse T cell–targeted TGF-β agonists on mouse CD4⁺ and CD8⁺ T cells *in vitro* and *in vivo*. **A.** Dose–response curves showing CD25, CD69, GITR, and T-bet expression in *in vitro*–cultured CD4⁺ T cells treated with the indicated proteins on day 3. **B.** Dose–response curves showing CD25, CD69, GITR, and T-bet expression, in *in vitro*–cultured CD8⁺ T cells treated with the indicated proteins on day 3. **C.** Quantification of the frequencies of TNF-α-producing cells among donor OT-II or OT-I cells in the DLNs of the indicated mice on day 7 after OVA immunization. Data in **A-C** are representative of two independent experiments. Data are presented as mean ± s.e.m. The statistics were obtained by one-way ANOVA coupled with Dunnett’s multiple-comparisons test **(C)**.

**Figure S3.**
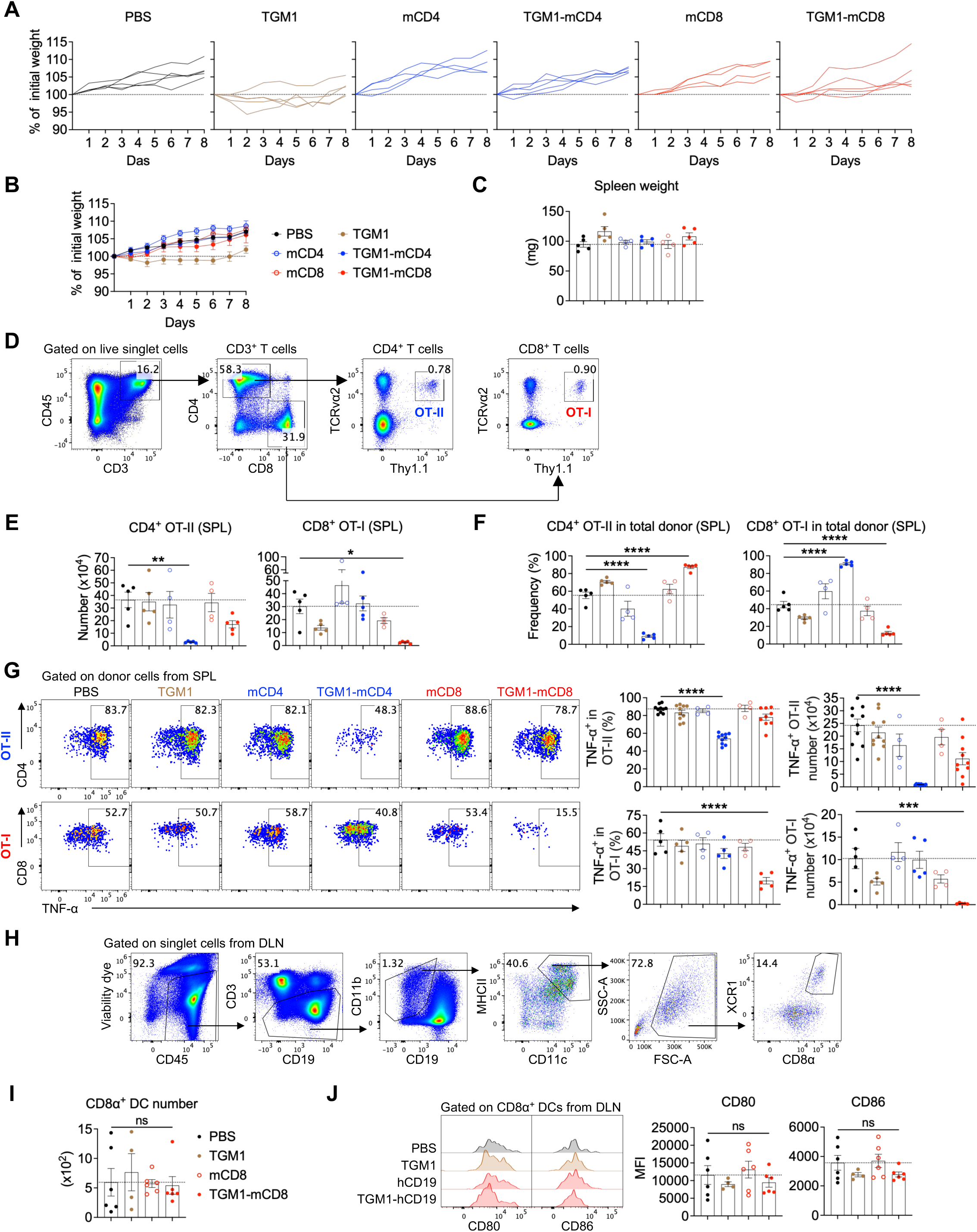
Mouse T cell-targeted TGF-β agonists silence antigen-specific T and B cell responses in OVA-immunized mice. **A.** Individual body weight change curves of mice from the indicated groups. **B.** Mean body weight change curves of mice from the indicated groups. **C.** Quantification of spleen weights of mice from the indicated groups. **D.** Gating strategy for donor CD4⁺ OT-II and CD8⁺ OT-I cells in DLNs. **E.** Quantification of donor OT-II and OT-I cell numbers in spleens of the indicated mice. **F.** Quantification of the proportion of OT-II or OT-I cells within total donor cells in spleens of the indicated mice. **G.** Representative flow cytometry plots and quantification of TNF-α–producing OT-II or OT-I cell frequencies and numbers in spleens of the indicated mice. **H.** Gating strategy for CD8α⁺ type 1 conventional dendritic cell (cDC1) in DLNs. **I.** Quantification of CD8α⁺ cDC1 cell numbers in DLNs of the indicated mice at ∼48 h after OVA immunization. **J.** Representative flow cytometry plots and quantification of CD80 and CD86 expression in CD8α⁺ cDC1 cells in DLNs of the indicated mice at ∼48 h after OVA immunization. Data in **A-J** are representative of two independent experiments. Data are presented as mean ± s.e.m. The statistics were obtained by one-way ANOVA coupled with Dunnett’s multiple-comparisons test **(E-G, I, and J)**.

**Figure S4.**
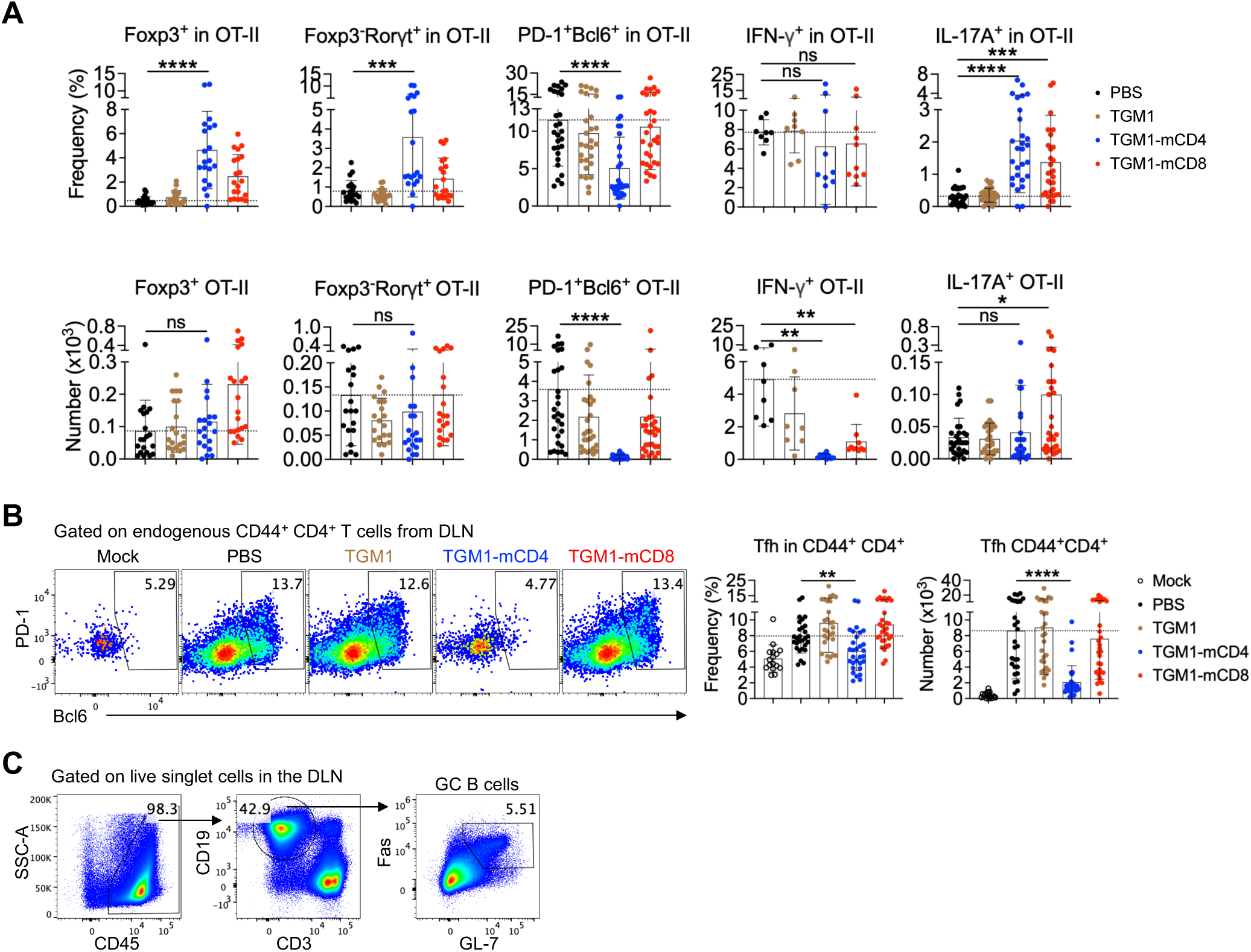
T cell phenotypes in OVA-immunized mice treated with mouse T cell–targeted TGF-β agonists. **A.** Quantification of Foxp3⁺, Foxp3⁻RORγt⁺, PD-1⁺Bcl6⁺, IFN-γ–producing, and IL-17A–producing OT-II cell frequencies (Upper panel) and numbers (Bottom panel) in the DLNs of mice from the indicated groups. **B.** Representative flow cytometry plots and quantification of PD-1⁺Bcl6⁺ Tfh cell frequencies and absolute numbers among endogenous CD4⁺ T cells in the DLNs of mice from the indicated groups. **C.** Gating strategy for GC B cells in DLNs. Data in **A-C** are pooled from two independent experiments. Data are presented as mean ± s.d. The statistics were obtained by one-way ANOVA coupled with Dunnett’s multiple-comparisons test **(A-C)**.

**Figure S5.**
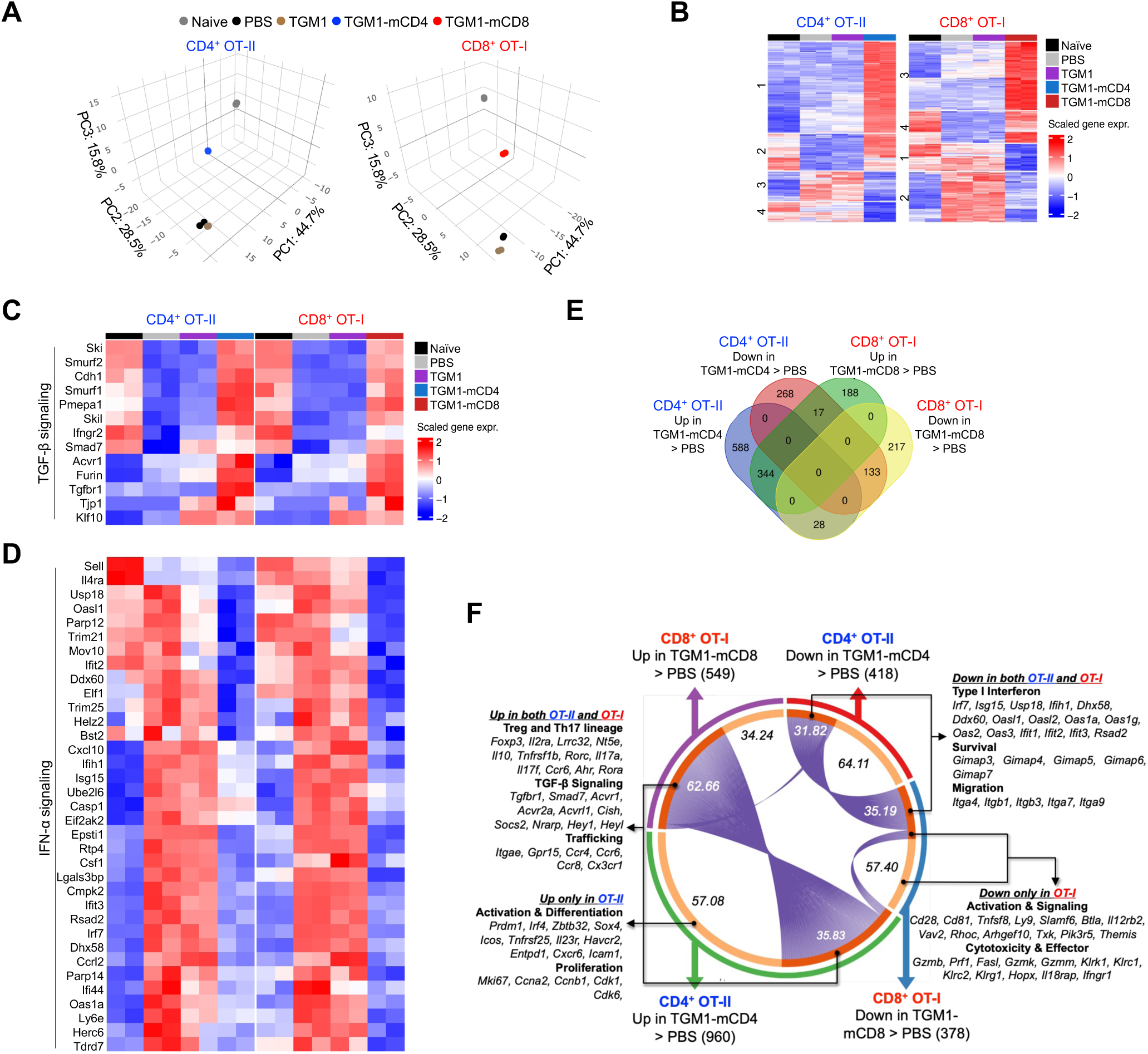
Transcriptional profiles of T cells from OVA-immunized mice treated with mouse T cell–targeted TGF-β agonists. **A.** Three-dimensional PCA plots showing the transcriptional profiles of OT-II or OT-I samples across the indicated groups. **B.** Gene expression heatmaps showing the transcriptional profiles of OT-II or OT-I samples across the indicated groups. **C.** Gene expression heatmaps showing expression levels of TGF-β signaling signature genes in OT-II or OT-I cells across the indicated groups. **D.** Gene expression heatmaps showing expression levels of IFN-α signaling signature genes in OT-II or OT-I cells across the indicated groups. **E.** Venn diagrams showing overlapping DEGs in OT-II and OT-I cells between T cell–targeted TGM1 and PBS groups. **F.** Chord diagram illustrating overlapping signature DEGs in OT-II and OT-I cells between T cell–targeted TGM1 and PBS groups.

**Figure S6.**
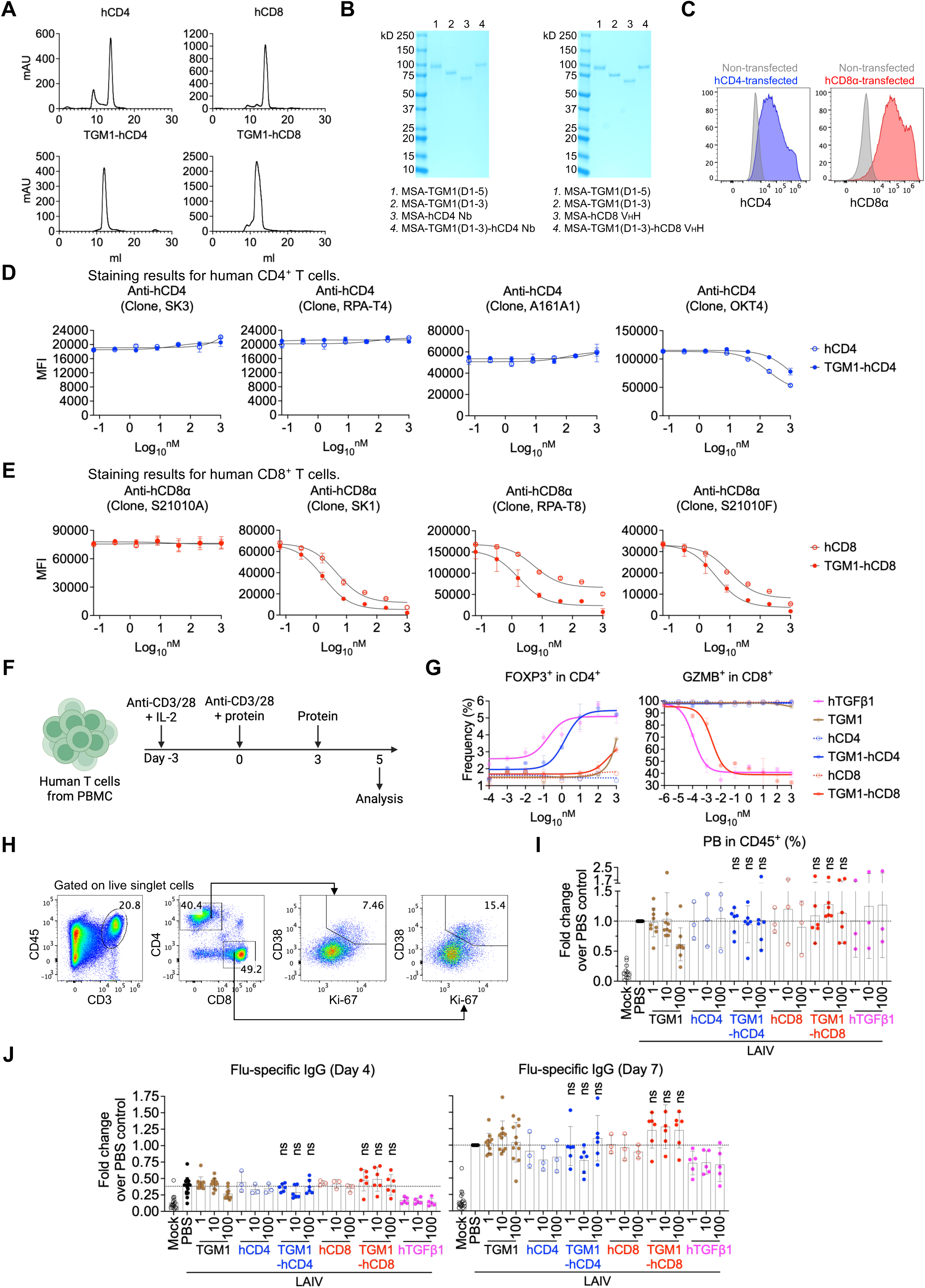
Design and characterization of human T cell-targeted TGF-β agonists. **A.** Analytical size-exclusion chromatography (SEC) profiles of the indicated proteins after Ni-NTA affinity purification. The major elution peak represents the purified protein of interest. **B.** SDS–PAGE analysis showing the molecular weights and purity of the indicated recombinant proteins. **C.** Representative flow cytometry plots showing human CD4 and CD8α expression in human CD4- or CD8α-transduced TGF-β reporter HEK293 cells. **D.** Dose–response curves showing staining intensities of the corresponding commercial antibodies on human PBMC CD4⁺ T cells after incubation with the indicated proteins. **E.** Dose–response curves showing staining intensities of the corresponding commercial antibodies on human PBMC CD8⁺ T cells after incubation with the indicated proteins. **F.** Experimental design of *in vitro* human PBMC T cell culture in the presence of the indicated proteins. **G.** Dose–response curves showing FOXP3⁺ cell frequencies in *in vitro*–cultured CD4⁺ T cells and Granzyme B–producing cell frequencies in *in vitro*–cultured CD8⁺ T cells treated with the indicated proteins on day 5. **H.** Gating strategy for CD38⁺Ki-67⁺ CD4⁺ and CD8⁺ T cells in *in vitro*–cultured human spleen organoids. **I.** Quantification of PB frequencies among total CD45⁺ leukocytes from the indicated groups. **J.** Quantification of flu-specific IgG levels in culture supernatants from the indicated groups on days 4 and 7. Data in **A-E and G** are representative of two independent experiments. Data in **I and J** are pooled from two independent experiments. Data in **D, E, and G** are presented as mean ± s.e.m. Data in **I and J** are presented as mean ± s.d. The statistics were obtained by one-way ANOVA coupled with Dunnett’s multiple-comparisons test **(I and J)**.

**Figure S7.**
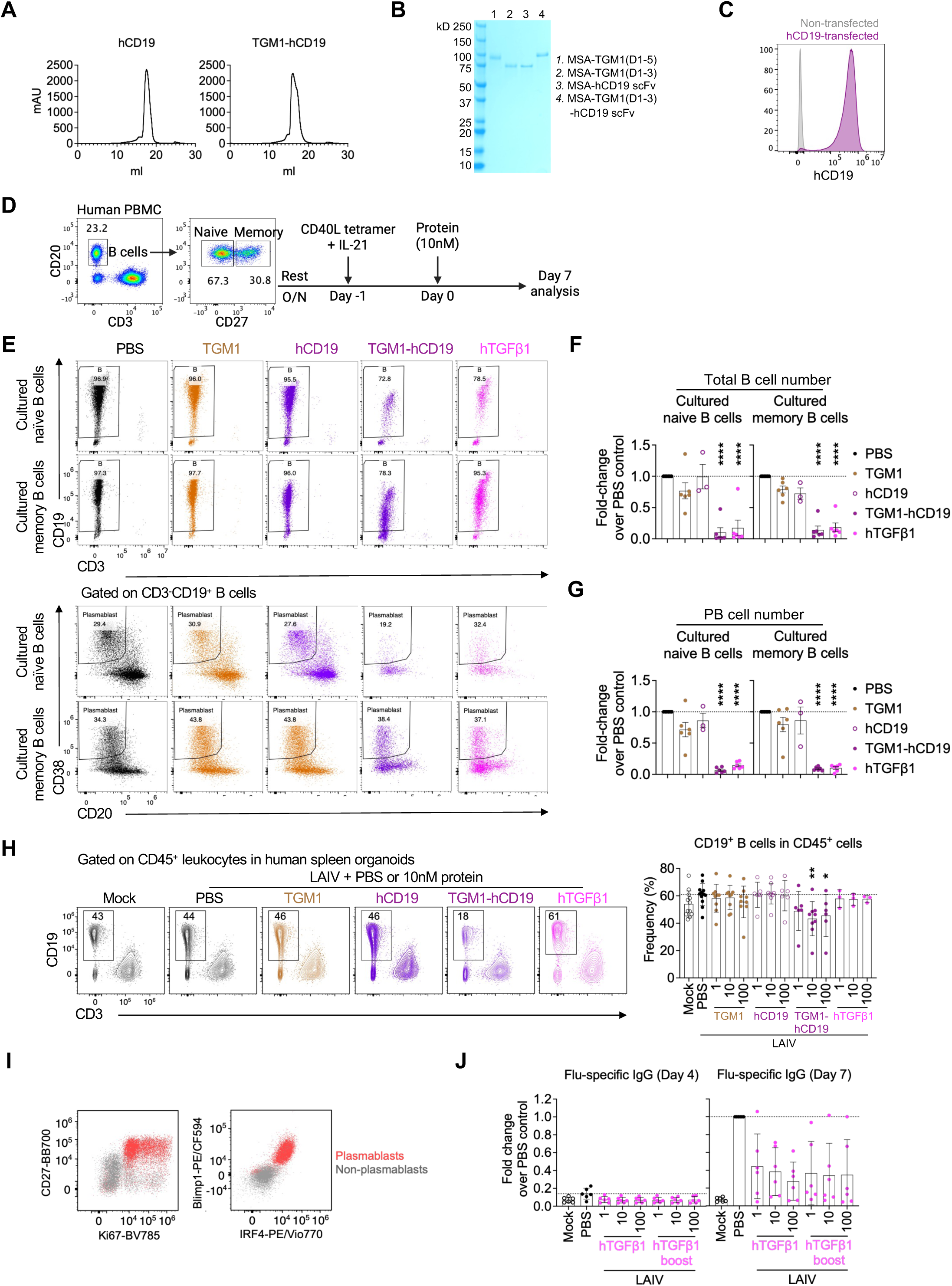
Design and characterization of human B cell-targeted TGF-β agonist. **A.** Analytical size-exclusion chromatography (SEC) profiles of the indicated proteins after Ni-NTA affinity purification. The major elution peak represents the purified protein of interest. **B.** SDS–PAGE analysis showing the molecular weight and purity of the indicated recombinant proteins. **C.** Representative flow cytometry plots showing human CD19 expression in CD19-transduced HEK293 cells. **D.** Schematic diagram illustrating sorting and *in vitro* culture of naïve and memory B cells isolated from human PBMCs. **E.** Representative flow cytometry plots of CD19 staining and PB differentiation in *in vitro*–cultured naïve and memory B cells. **F.** Quantification of normalized absolute counts of total B cells across indicated groups. Data include four healthy controls and two SLE patients. **G.** Quantification of normalized absolute counts of PB cells across indicated groups. Data include four healthy controls and two SLE patients. **H.** Representative flow cytometry plots of CD19 staining in human spleen organoids and quantification of B cell frequencies among total CD45⁺ leukocytes across indicated groups. **I.** Representative flow cytometry plots of Ki67, CD27, IRF4, and BLIMP1 expression in PBs (CD3⁻CD19⁺CD20^dim^CD38^high^) and non-PB populations in a human spleen organoid. **J.** Flu-specific IgG in culture supernatants at days 4 and 7, measured by ELISA and expressed as fold change relative to the day-7 LAIV-alone condition of the same donor (n = 6 donors). “boost” denotes re-administration of rhTGF-β1 at the same concentration during the day-4 medium change. Data in **A-G** are representative of two independent experiments. Data in **H-J** are pooled from three independent experiments. Data in **F and G** are presented as mean ± s.e.m. Data in **H and J** are presented as mean ± s.d. The statistics were obtained by one-way ANOVA coupled with Dunnett’s multiple-comparisons test **(F-H)**.

**Figure S8.**
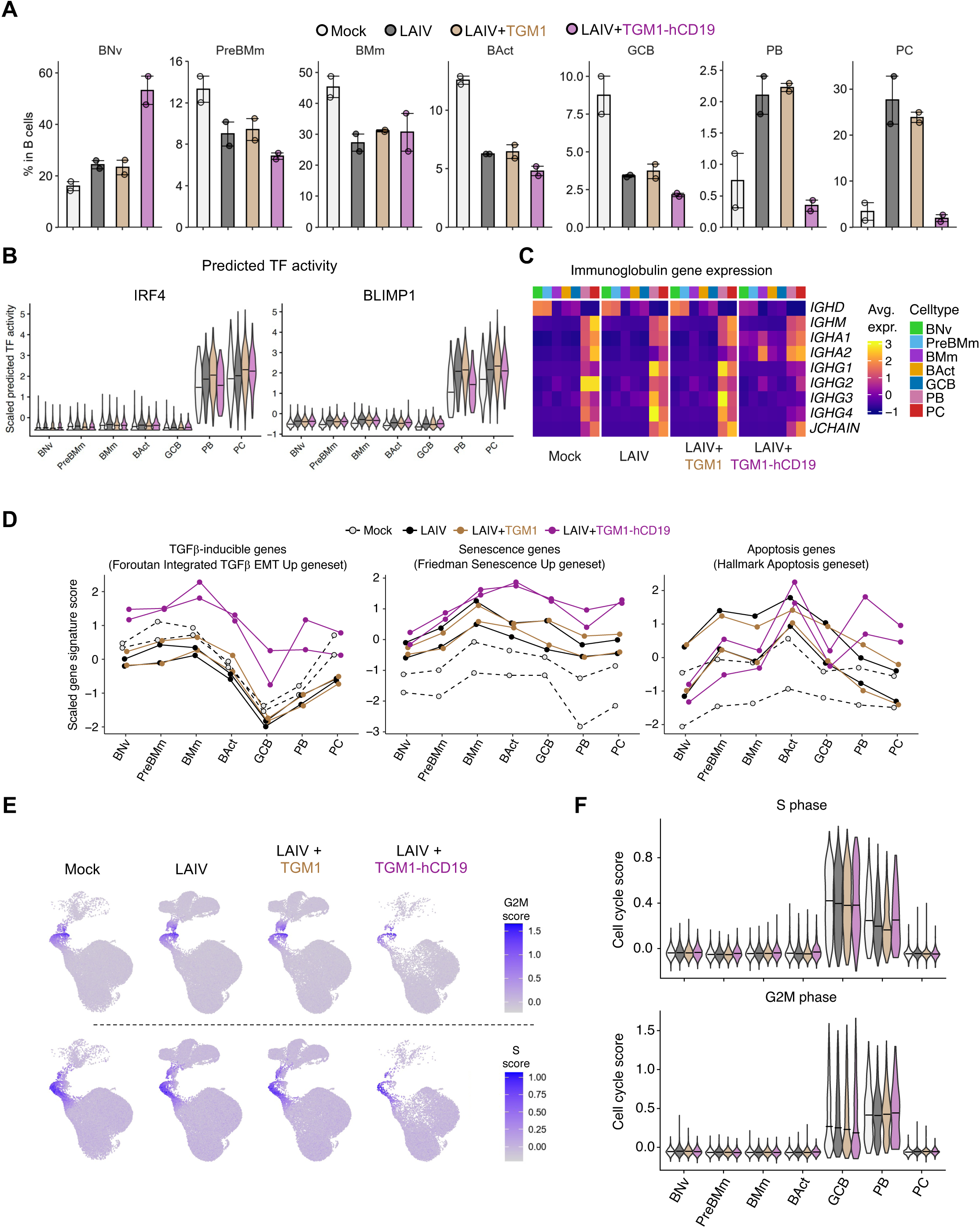
scRNA-seq analysis of B cells in human spleen organoids treated with a human B cell–targeted TGF-β agonist. **A**. Quantification of frequencies of distinct B cell subsets among total B cells, derived from scRNA-seq data across indicated groups. **B.** SCENIC-based prediction of IRF4 and BLIMP1 transcription factor activity across distinct B cell clusters in indicated groups. **C.** Heatmaps showing scaled average expression of immunoglobulin genes across distinct B cell clusters in indicated groups. **D.** Scaled signature scores for TGF-β–inducible, senescence-associated, and apoptosis-related gene sets, aggregated by sample. **E.** UMAP plots showing cell-cycle gene signature scores across indicated groups. **F.** Quantification of cell-cycle gene signature scores across distinct B cell clusters in indicated groups.

**Figure S9.**
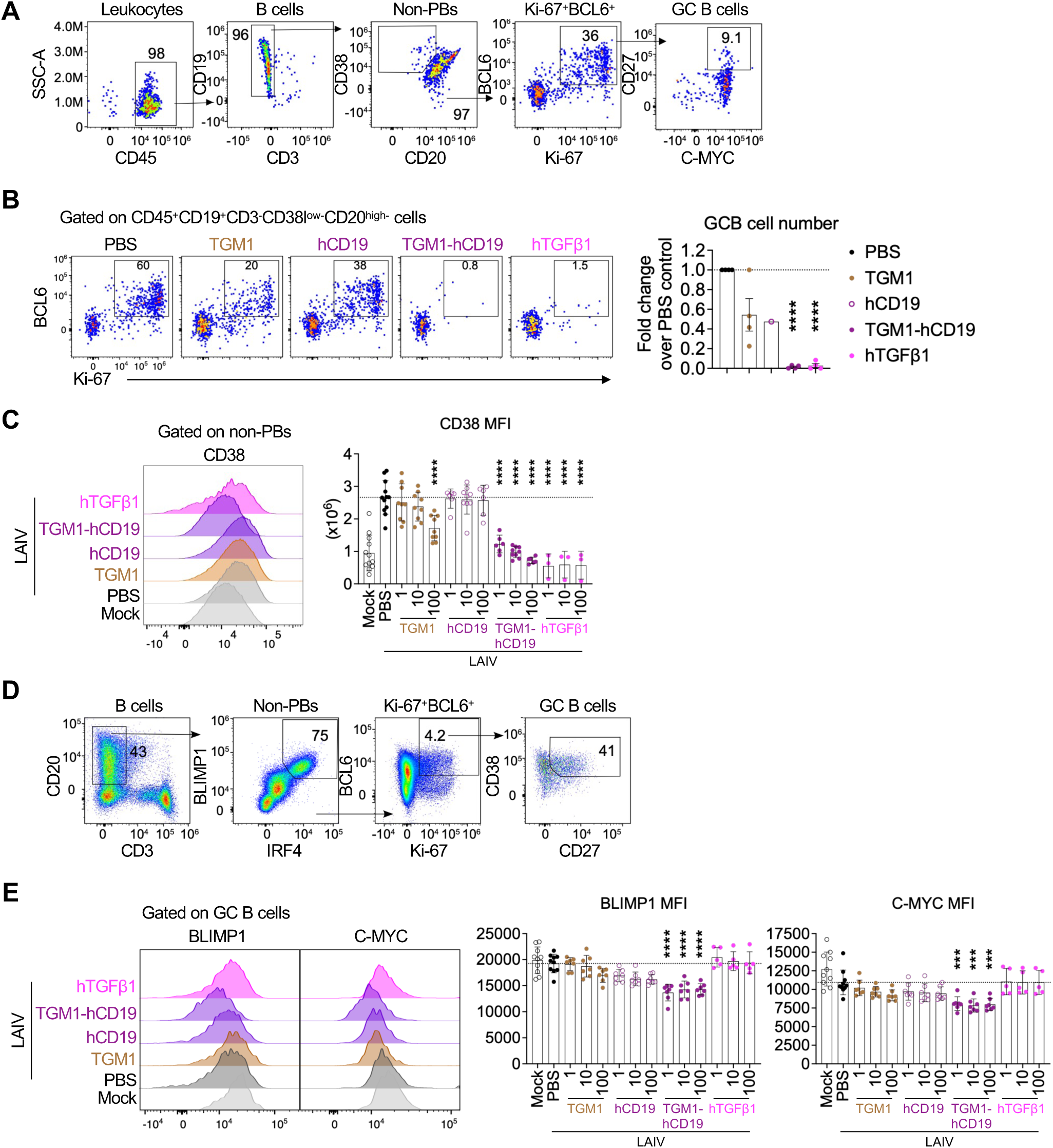
Suppressive effects of a human B cell–targeted TGF-β agonist on antigen-driven B cell differentiation in human spleen organoids. **A.** Gating strategy for GC B cells derived from *in vitro*–cultured, sort-purified naïve B cells stimulated with CD40L and IL-21 (see **Fig. S7C**). **B.** Representative flow cytometry plots and quantification of GC B cell frequencies and normalized absolute GC B cell counts across indicated groups. **C.** Representative flow cytometry plots and quantification of CD38 expression in the non-PBs in the human spleen organoid LAIV assay across indicated groups. **D.** Gating strategy for GC B cells in human spleen organoids. **E.** Representative flow cytometry plots and quantification of BLIMP1 and c-MYC expression in GC B cells from human spleen organoids across indicated groups. Data in **A and B** are representative of two independent experiments. Data in **C-E** are pooled from three independent experiments. Data in **B** are presented as mean ± s.e.m. Data in **C and E** are presented as mean ± s.d. The statistics were obtained by one-way ANOVA coupled with Dunnett’s multiple-comparisons test **(B, C, and E)**.

**Figure S10.**
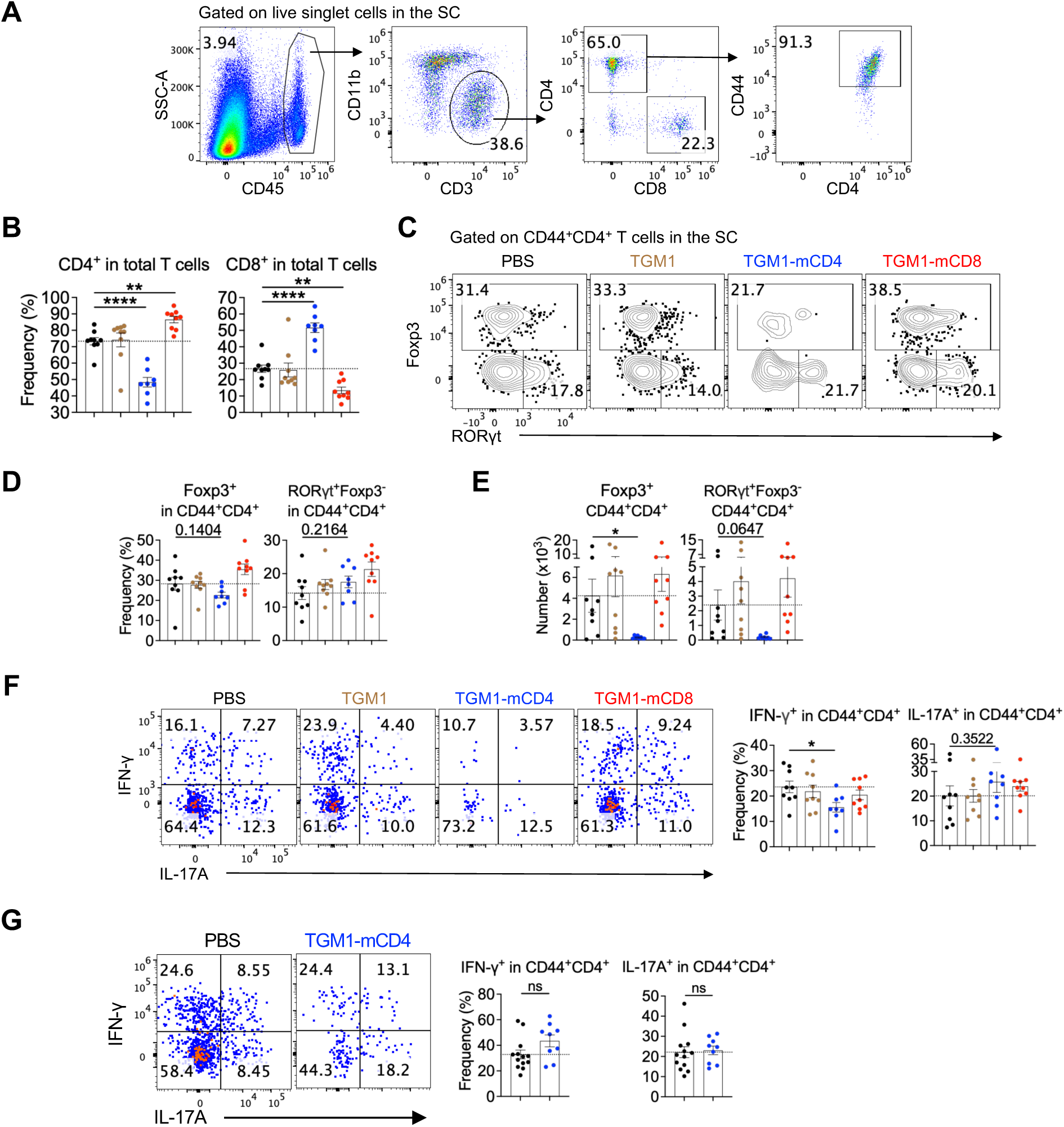
Therapeutic efficacy of a mouse T cell–targeted TGF-β agonist in the MOG₃₅–₅₅-induced EAE model. **A.** Gating strategy for CD4⁺ and CD8⁺ T cells from SCs. **B.** Quantification of CD4⁺ and CD8⁺ T cell proportions among total T cells in SCs of mice from the indicated groups. **C.** Representative flow cytometry plots showing Foxp3⁺ and RORγt⁺ cell frequencies within CD44⁺CD4⁺ T cells in the SCs of mice from the indicated groups. **D.** Quantification of Foxp3⁺ and RORγt⁺ cell frequencies within CD44⁺CD4⁺ T cells in the SCs of mice from the indicated groups. **E.** Quantification of Foxp3⁺ and RORγt⁺ CD44⁺CD4⁺ T cell numbers in the SCs of mice from the indicated groups. **F.** Representative flow cytometry plots and quantification of IFN-γ– and IL-17A–producing cell frequencies within CD44⁺CD4⁺ T cells in the SCs of mice from the indicated groups. **G.** Representative flow cytometry plots and quantification of the frequencies of IFN-γ- and IL-17A-producing cells within CD44⁺CD4⁺ T cells in the SCs. Data in **B-G** are representative of two independent experiments. Data are presented as mean ± s.e.m. The statistics were obtained by one-way ANOVA coupled with Dunnett’s multiple-comparisons test **(B-F)** or unpaired Welch’s t-test (two-tailed) **(G)**.

**Figure S11.**
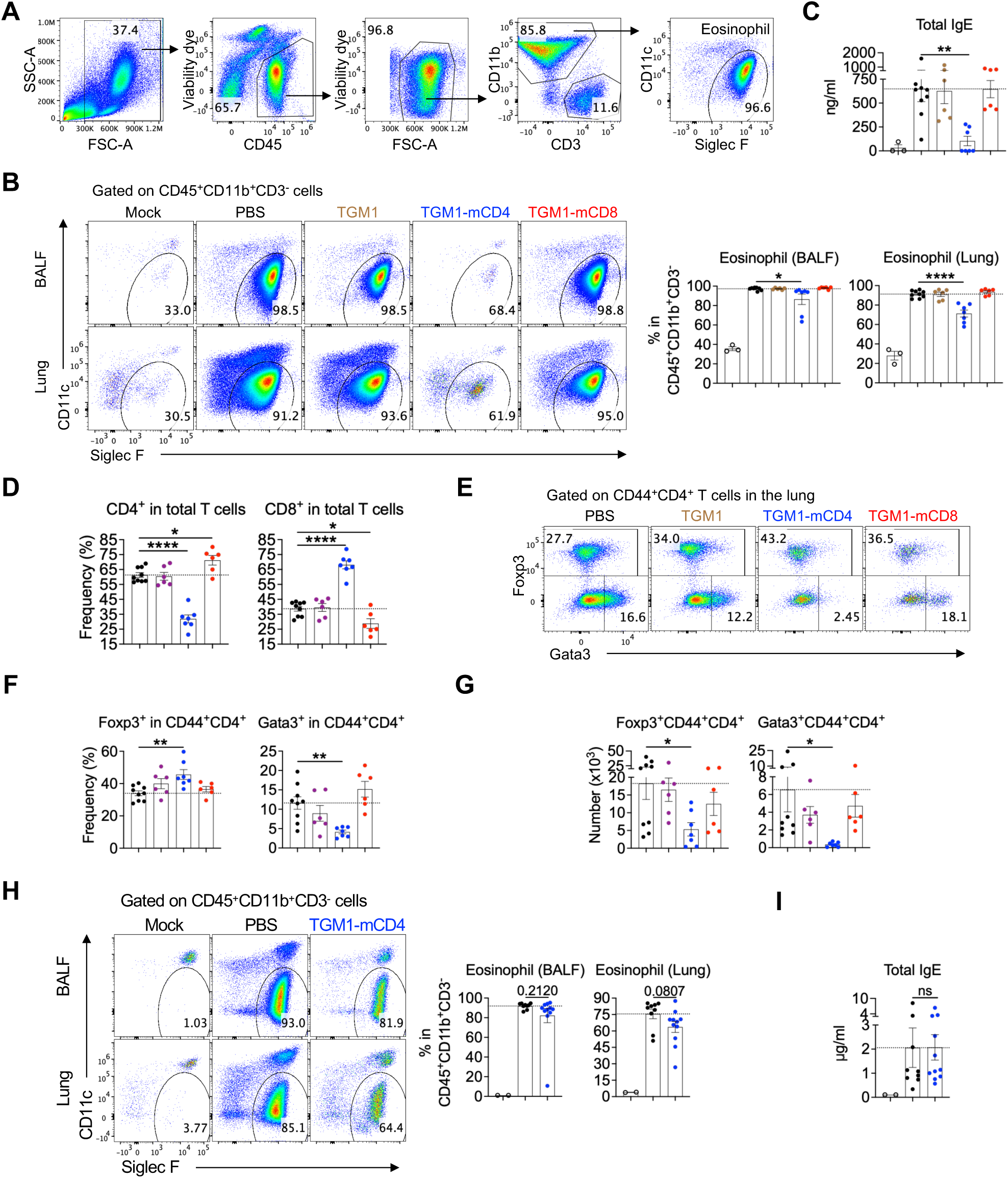
Therapeutic efficacy of a mouse T cell–targeted TGF-β agonist in the OVA-induced asthma model. **A.** Gating strategy for eosinophils in BALF and lung tissue. **B.** Representative flow cytometry plots and quantification of eosinophils among CD45⁺CD11b⁺CD3⁻ cells in BALF and lungs of mice from the indicated groups. **C.** Quantification of serum total IgE levels in mice from the indicated groups. **D.** Quantification of CD4⁺ and CD8⁺ T cell proportions among total T cells in lungs of mice from the indicated groups. **E.** Representative flow cytometry plots of Foxp3⁺ and Gata3⁺ cell frequencies within CD44⁺CD4⁺ T cells in the lungs of mice from the indicated groups. **F.** Quantification of Foxp3⁺ and Gata3⁺ cell frequencies within CD44⁺CD4⁺ T cells in the lungs of mice from the indicated groups. **G.** Quantification of Foxp3⁺ and Gata3⁺ CD44⁺CD4⁺ T cell numbers in the lungs of mice from the indicated groups. **H.** Representative flow cytometry plots and quantification of eosinophils among CD45⁺CD11b⁺CD3⁻ cells in BALF and lungs of mice from the indicated groups. **I.** Quantification of serum total IgE levels in mice from the indicated groups. Data in **B-I** are representative of two independent experiments. Data are presented as mean ± s.e.m. The statistics were obtained by one-way ANOVA coupled with Dunnett’s multiple-comparisons test **(B-G)** or unpaired Welch’s t-test (two-tailed) **(H and I)**.

**Figure S12.**
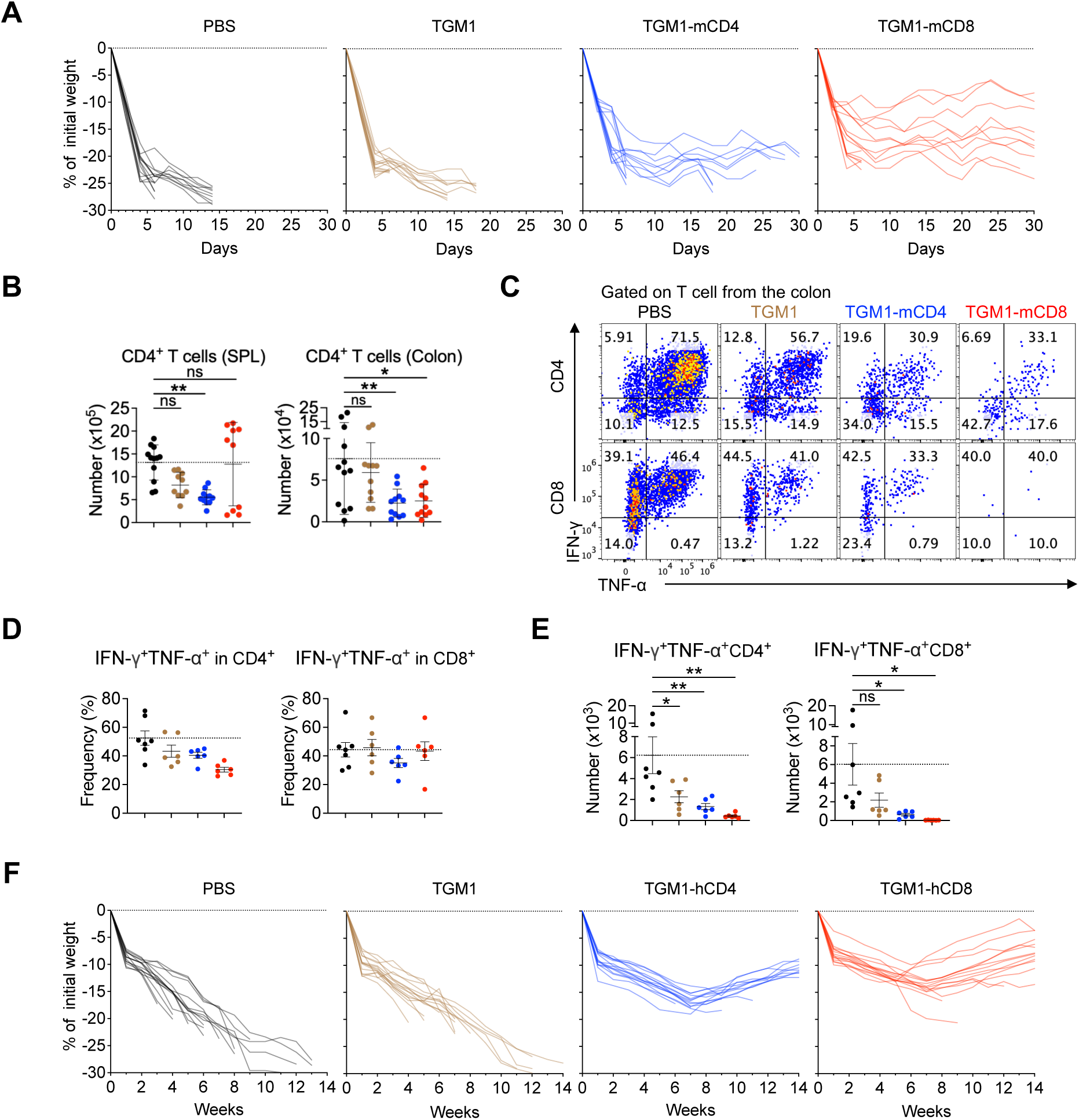
Therapeutic efficacy of T cell–targeted TGF-β agonists in mouse and human GVHD models. **A.** Individual body weight change curves of mice from the indicated groups. **B.** Quantification of CD4⁺ T cell numbers in spleen and colon of mice from the indicated groups. **C.** Representative flow cytometry plots showing IFN-γ– and TNF-α–producing cell frequencies within CD4⁺ and CD8⁺ T cells in the colons of mice from the indicated groups. **D.** Quantification of IFN-γ– and TNF-α–producing cell frequencies within CD4⁺ and CD8⁺ T cells in the colons of mice from the indicated groups. **E.** Quantification of IFN-γ– and TNF-α–producing CD4⁺ and CD8⁺ T cell numbers in the colons of mice from the indicated groups. **F.** Individual body weight change curves of mice from the indicated groups. Data in **A, B and F** are pooled from two independent experiments. Data in **C-E** are representative of two independent experiments. Data in **B** are presented as mean ± s.d. Data in **D and E** are presented as mean ± s.e.m. The statistics were obtained by one-way ANOVA coupled with Dunnett’s multiple-comparisons test **(B and E)**.

**Supplementary Table 1.** Assessment of clinical GVHD in transplanted mice.

| **Mouse allogeneic GVHD model** |  |  |  |  |  |
| --- | --- | --- | --- | --- | --- |
| Criteria | Grade 0 | Grade 0.5 | Grade 1.0 | Grade 1.5 | Grade 2.0 |
| Weight loss | <10% |  | >10% to <25% |  | >25% |
| Posture | No hunch | Slight hunch, straightens when walks | Animal stays hunched when walks | Animal does not straighten out | Animal tends to stand on rear toes |
| Mobility | Very mobile, hard to catch | Slower than naïve, easier to catch | Not moving, but will move when poked | Not moving, moves slightly when poked | Not moving, won't move if poked |
| Fur | No fur pathology | Ridging on the side of belly or nape of neck | Ridging across or the side of belly + neck | Unkempt matted and ruffled fur | Badly matted fur on belly, and on top |
| Skin | No redness, abrasions, lesions or scaling present | Redness in one area only | Abrasions in 1 area, or mild abrasions in 2 areas | Bad abrasions in 2 areas | Extremely bad abrasion, cracking skin, dried blood etc. |
| **Humanized xenogeneic GVHD model** |  |  |  |  |  |
| Criteria | Grade 0 | Grade 1 |  | Grade 2 |  |
| Weight loss | < 10% | >10% to <25% |  | > 25% |  |
| Posture | Normal | Hunching noted only at rest |  | Severe hunching impairs movement |  |
| Activity | Normal | Mild to moderately decreased |  | Stationary unless stimulated |  |
| Fur texture | Normal | Mild to moderate ruffling |  | Sever ruffling poor grooming |  |
| Skin Integrity | Normal | Scaling of paws and tail |  | Obvious areas of denuded skin |  |

## Reference

1 Sun, L., Su, Y., Jiao, A., Wang, X. & Zhang, B. T cells in health and disease. Signal Transduct Target Ther 8, 235 (2023). 10.1038/s41392-023-01471-y

2 Martin, F. & Chan, A. C. Pathogenic Roles of B Cells in Human Autoimmunity: Insights from the Clinic. Immunity 20, 517–527 (2004). 10.1016/S1074-7613(04)00112-8

3 Edner, N. M., Carlesso, G., Rush, J. S. & Walker, L. S. K. Targeting co-stimulatory molecules in autoimmune disease. Nat Rev Drug Discov 19, 860–883 (2020). 10.1038/s41573-020-0081-9

4 Chung, J. B., Brudno, J. N., Borie, D. & Kochenderfer, J. N. Chimeric antigen receptor T cell therapy for autoimmune disease. Nat Rev Immunol 24, 830–845 (2024). 10.1038/s41577-024-01035-3

5 Chan, A. C. & Carter, P. J. Therapeutic antibodies for autoimmunity and inflammation. Nat Rev Immunol 10, 301–316 (2010). 10.1038/nri2761

6 Jacobson, P., Uberti, J., Davis, W. & Ratanatharathorn, V. Tacrolimus: a new agent for the prevention of graft-versus-host disease in hematopoietic stem cell transplantation. Bone Marrow Transplantation 22, 217–225 (1998). 10.1038/sj.bmt.1701331

7 Choi, S. W. & Reddy, P. Current and emerging strategies for the prevention of graft-versus-host disease. Nat Rev Clin Oncol 11, 536–547 (2014). 10.1038/nrclinonc.2014.102

8 Herold, K. C. et al. An Anti-CD3 Antibody, Teplizumab, in Relatives at Risk for Type 1 Diabetes. N Engl J Med 381, 603–613 (2019). 10.1056/NEJMoa1902226

9 Aronson, R. et al. Low-dose otelixizumab anti-CD3 monoclonal antibody DEFEND-1 study: results of the randomized phase III study in recent-onset human type 1 diabetes. Diabetes Care 37, 2746–2754 (2014). 10.2337/dc13-0327

10 Kappos, L. et al. Ocrelizumab in relapsing-remitting multiple sclerosis: a phase 2, randomised, placebo-controlled, multicentre trial. Lancet 378, 1779–1787 (2011). 10.1016/S0140-6736(11)61649-8

11 Montalban, X. et al. Ocrelizumab versus Placebo in Primary Progressive Multiple Sclerosis. N Engl J Med 376, 209–220 (2017). 10.1056/NEJMoa1606468

12 Furie, R. A. et al. Efficacy and Safety of Obinutuzumab in Active Lupus Nephritis. N Engl J Med 392, 1471–1483 (2025). 10.1056/NEJMoa2410965

13 Muller, F. et al. CD19 CAR T-Cell Therapy in Autoimmune Disease - A Case Series with Follow-up. N Engl J Med 390, 687–700 (2024). 10.1056/NEJMoa2308917

14 Muller, F. et al. CD19 CAR-T cells for treatment-refractory autoimmune diseases: the phase 1/2 CASTLE basket trial. Nat Med 32, 1142–1151 (2026). 10.1038/s41591-025-04185-6

15 Ramos, E. L. et al. Teplizumab and beta-Cell Function in Newly Diagnosed Type 1 Diabetes. N Engl J Med 389, 2151–2161 (2023). 10.1056/NEJMoa2308743

16 Sutherland, N. M. et al. Association of CD19(+)-targeted chimeric antigen receptor (CAR) T-cell therapy with hypogammaglobulinemia, infection, and mortality. J Allergy Clin Immunol 155, 605–615 (2025). 10.1016/j.jaci.2024.10.021

17 Brudno, J. N. & Kochenderfer, J. N. Current understanding and management of CAR T cell-associated toxicities. Nat Rev Clin Oncol 21, 501–521 (2024). 10.1038/s41571-024-00903-0

18 Hansel, T. T., Kropshofer, H., Singer, T., Mitchell, J. A. & George, A. J. The safety and side effects of monoclonal antibodies. Nat Rev Drug Discov 9, 325–338 (2010). 10.1038/nrd3003

19 Li, M. O., Wan, Y. Y., Sanjabi, S., Robertson, A. K. & Flavell, R. A. Transforming growth factor-beta regulation of immune responses. Annu Rev Immunol 24, 99–146 (2006). 10.1146/annurev.immunol.24.021605.090737

20 Chen, W. TGF-beta Regulation of T Cells. Annu Rev Immunol 41, 483–512 (2023). 10.1146/annurev-immunol-101921-045939

21 Chen, W. et al. Conversion of peripheral CD4+CD25-naive T cells to CD4+CD25+ regulatory T cells by TGF-beta induction of transcription factor Foxp3. J Exp Med 198, 1875–1886 (2003). 10.1084/jem.20030152

22 Bettelli, E. et al. Reciprocal developmental pathways for the generation of pathogenic effector TH17 and regulatory T cells. Nature 441, 235–238 (2006). 10.1038/nature04753

23 Mangan, P. R. et al. Transforming growth factor-beta induces development of the T(H)17 lineage. Nature 441, 231–234 (2006). 10.1038/nature04754

24 Zhang, N. & Bevan, M. J. Transforming growth factor-beta signaling controls the formation and maintenance of gut-resident memory T cells by regulating migration and retention. Immunity 39, 687–696 (2013). 10.1016/j.immuni.2013.08.019

25 Cazac, B. B. & Roes, J. TGF-β Receptor Controls B Cell Responsiveness and Induction of IgA In Vivo. Immunity 13, 443–451 (2000). 10.1016/S1074-7613(00)00044-3

26 Gorelik, L. & Flavell, R. A. Abrogation of TGFβ Signaling in T Cells Leads to Spontaneous T Cell Differentiation and Autoimmune Disease. Immunity 12, 171–181 (2000). 10.1016/S1074-7613(00)80170-3

27 Shull, M. M. et al. Targeted disruption of the mouse transforming growth factor-β1 gene results in multifocal inflammatory disease. Nature 359, 693–699 (1992). 10.1038/359693a0

28 Kulkarni, A. B. et al. Transforming growth factor beta 1 null mutation in mice causes excessive inflammatory response and early death. Proceedings of the National Academy of Sciences 90, 770–774 (1993). doi:10.1073/pnas.90.2.770

29 Massague, J. & Sheppard, D. TGF-beta signaling in health and disease. Cell 186, 4007–4037 (2023). 10.1016/j.cell.2023.07.036

30 Lambert, F. et al. Antibody-mediated TGF-beta1 activation for the treatment of diseases caused by deleterious T cell activity. Cell Rep 44, 116061 (2025). 10.1016/j.celrep.2025.116061

31 Johnston, C. J. C. et al. A structurally distinct TGF-beta mimic from an intestinal helminth parasite potently induces regulatory T cells. Nat Commun 8, 1741 (2017). 10.1038/s41467-017-01886-6

32 Mukundan, A. et al. Convergent evolution of a parasite-encoded complement control protein-scaffold to mimic binding of mammalian TGF-beta to its receptors, TbetaRI and TbetaRII. J Biol Chem 298, 101994 (2022). 10.1016/j.jbc.2022.101994

33 Sun, Q. et al. Facile induction of immune tolerance by an interleukin-2-TGFbeta surrogate agonist. Nature (2026). 10.1038/s41586-026-10208-0

34 van Dinther, M. et al. CD44 acts as a coreceptor for cell-specific enhancement of signaling and regulatory T cell induction by TGM1, a parasite TGF-beta mimic. Proc Natl Acad Sci U S A 120, e2302370120 (2023). 10.1073/pnas.2302370120

35 Cobbold, S. P., Jayasuriya, A., Nash, A., Prospero, T. D. & Waldmann, H. Therapy with monoclonal antibodies by elimination of T-cell subsets in vivo. Nature 312, 548–551 (1984). 10.1038/312548a0

36 Rashidian, M. et al. Predicting the response to CTLA-4 blockade by longitudinal noninvasive monitoring of CD8 T cells. J Exp Med 214, 2243–2255 (2017). 10.1084/jem.20161950

37 Mildner, A. & Jung, S. Development and function of dendritic cell subsets. Immunity 40, 642–656 (2014). 10.1016/j.immuni.2014.04.016

38 Hirai, T. et al. Competition for Active TGFbeta Cytokine Allows for Selective Retention of Antigen-Specific Tissue-Resident Memory T Cells in the Epidermal Niche. Immunity 54, 84–98 e85 (2021). 10.1016/j.immuni.2020.10.022

39 Topchyan, P., Lin, S. & Cui, W. The Role of CD4 T Cell Help in CD8 T Cell Differentiation and Function During Chronic Infection and Cancer. Immune Netw 23, e41 (2023). 10.4110/in.2023.23.e41

40 Traenkle, B. et al. Single-Domain Antibodies for Targeting, Detection, and In Vivo Imaging of Human CD4(+) Cells. *Front Immunol* **12**, 799910 (2021). 10.3389/fimmu.2021.799910

41 Sriraman, S. K. et al. Development of an (18)F-labeled anti-human CD8 VHH for same-day immunoPET imaging. Eur J Nucl Med Mol Imaging 50, 679–691 (2023). 10.1007/s00259-022-05998-0

42 Wagar, L. E. et al. Modeling human adaptive immune responses with tonsil organoids. Nat Med 27, 125–135 (2021). 10.1038/s41591-020-01145-0

43 Mallajosyula, V. et al. Coupling antigens from multiple subtypes of influenza can broaden antibody and T cell responses. Science 386, 1389–1395 (2024). doi:10.1126/science.adi2396

44 He, C. et al. CD19 CAR antigen engagement mechanisms and affinity tuning. Science Immunology 8, eadf1426 (2023). doi:10.1126/sciimmunol.adf1426

45 Nicholson, I. C. et al. Construction and characterisation of a functional CD19 specific single chain Fv fragment for immunotherapy of B lineage leukaemia and lymphoma. Molecular Immunology 34, 1157–1165 (1997). 10.1016/S0161-5890(97)00144-2

46 Massoni-Badosa, R. et al. An atlas of cells in the human tonsil. Immunity 57, 379–399 e318 (2024). 10.1016/j.immuni.2024.01.006

47 Aibar, S. et al. SCENIC: single-cell regulatory network inference and clustering. Nat Methods 14, 1083–1086 (2017). 10.1038/nmeth.4463

48 La Manno, G. et al. RNA velocity of single cells. Nature 560, 494–498 (2018). 10.1038/s41586-018-0414-6

49 Conter, L. et al. BLIMP1 controls GC B cell expansion and exit through regulating cell cycle progression and key transcription factors BCL6 and IRF4. Cell Rep 44, 115977 (2025). 10.1016/j.celrep.2025.115977

50 Finkin, S., Hartweger, H., Oliveira, T. Y., Kara, E. E. & Nussenzweig, M. C. Protein Amounts of the MYC Transcription Factor Determine Germinal Center B Cell Division Capacity. Immunity 51, 324–336 e325 (2019). 10.1016/j.immuni.2019.06.013

51 Johns, L. D., Flanders, K. C., Ranges, G. E. & Sriram, S. Successful treatment of experimental allergic encephalomyelitis with transforming growth factor-beta 1. The Journal of Immunology 147, 1792–1796 (1991). 10.4049/jimmunol.147.6.1792

52 Racke, M. K. et al. Prevention and treatment of chronic relapsing experimental allergic encephalomyelitis by transforming growth factor-beta 1. The Journal of Immunology 146, 3012–3017 (1991). 10.4049/jimmunol.146.9.3012

53 Santambrogio, L. et al. Studies on the mechanisms by which transforming growth factor-beta (TGF-beta) protects against allergic encephalomyelitis. Antagonism between TGF-beta and tumor necrosis factor. The Journal of Immunology 151, 1116–1127 (1993). 10.4049/jimmunol.151.2.1116

54 Kuruvilla, A. P. et al. Protective effect of transforming growth factor beta 1 on experimental autoimmune diseases in mice. Proceedings of the National Academy of Sciences 88, 2918–2921 (1991). doi:10.1073/pnas.88.7.2918

55 McGeachy, M. J. et al. TGF-beta and IL-6 drive the production of IL-17 and IL-10 by T cells and restrain T(H)-17 cell-mediated pathology. Nat Immunol 8, 1390–1397 (2007). 10.1038/ni1539

56 Veldhoen, M., Hocking, R. J., Flavell, R. A. & Stockinger, B. Signals mediated by transforming growth factor-beta initiate autoimmune encephalomyelitis, but chronic inflammation is needed to sustain disease. Nat Immunol 7, 1151–1156 (2006). 10.1038/ni1391

57 Melton, A. C. et al. Expression of alphavbeta8 integrin on dendritic cells regulates Th17 cell development and experimental autoimmune encephalomyelitis in mice. J Clin Invest 120, 4436–4444 (2010). 10.1172/JCI43786

58 Banovic, T. et al. TGF-beta in allogeneic stem cell transplantation: friend or foe? Blood 106, 2206–2214 (2005). 10.1182/blood-2005-01-0062

59 Chang, Y. et al. TGF-β specifies TFH versus TH17 cell fates in murine CD4+ T cells through c-Maf. Science Immunology 9, eadd4818 (2024). doi:10.1126/sciimmunol.add4818

60 Chaurio, R. A. et al. TGF-beta-mediated silencing of genomic organizer SATB1 promotes Tfh cell differentiation and formation of intra-tumoral tertiary lymphoid structures. Immunity 55, 115–128 e119 (2022). 10.1016/j.immuni.2021.12.007

61 Marshall, H. D. et al. The transforming growth factor beta signaling pathway is critical for the formation of CD4 T follicular helper cells and isotype-switched antibody responses in the lung mucosa. Elife 4, e04851 (2015). 10.7554/eLife.04851

62 Schmitt, N. et al. The cytokine TGF-beta co-opts signaling via STAT3-STAT4 to promote the differentiation of human TFH cells. Nat Immunol 15, 856–865 (2014). 10.1038/ni.2947

63 McCarron, M. J. & Marie, J. C. TGF-beta prevents T follicular helper cell accumulation and B cell autoreactivity. J Clin Invest 124, 4375–4386 (2014). 10.1172/JCI76179

64 Cohn, L., Homer, R. J., Marinov, A., Rankin, J. & Bottomly, K. Induction of Airway Mucus Production By T Helper 2 (Th2) Cells: A Critical Role For Interleukin 4 In Cell Recruitment But Not Mucus Production. Journal of Experimental Medicine 186, 1737–1747 (1997). 10.1084/jem.186.10.1737

65 Vinuesa, C. G., Linterman, M. A., Yu, D. & MacLennan, I. C. Follicular Helper T Cells. Annu Rev Immunol 34, 335–368 (2016). 10.1146/annurev-immunol-041015-055605

66 Victora, G. D. & Nussenzweig, M. C. Germinal Centers. Annu Rev Immunol 40, 413–442 (2022). 10.1146/annurev-immunol-120419-022408

67 Kato, A. & Kita, H. The immunology of asthma and chronic rhinosinusitis. Nat Rev Immunol 25, 569–587 (2025). 10.1038/s41577-025-01159-0

68 Li, M. O., Sanjabi, S. & Flavell, R. A. Transforming growth factor-beta controls development, homeostasis, and tolerance of T cells by regulatory T cell-dependent and -independent mechanisms. Immunity 25, 455–471 (2006). 10.1016/j.immuni.2006.07.011

69 Moreau, J. M., Velegraki, M., Bolyard, C., Rosenblum, M. D. & Li, Z. Transforming growth factor–β1 in regulatory T cell biology. Science Immunology 7, eabi4613 (2022). doi:10.1126/sciimmunol.abi4613

70 Yang, X. O. et al. Molecular antagonism and plasticity of regulatory and inflammatory T cell programs. Immunity 29, 44–56 (2008). 10.1016/j.immuni.2008.05.007

71 Chung, Y. et al. Critical regulation of early Th17 cell differentiation by interleukin-1 signaling. Immunity 30, 576–587 (2009). 10.1016/j.immuni.2009.02.007

72 Schoenberger, S. P., Toes, R. E. M., van der Voort, E. I. H., Offringa, R. & Melief, C. J. M. T-cell help for cytotoxic T lymphocytes is mediated by CD40–CD40L interactions. Nature 393, 480–483 (1998). 10.1038/31002

73 Bennett, S. R. M. et al. Help for cytotoxic-T-cell responses is mediated by CD40 signalling. Nature 393, 478–480 (1998). 10.1038/30996

74 Zhao, Z. et al. Dynamic expression of IL-21 in effector CD8 T cells helps control chronic infections. Science Immunology 11, eaec5171 (2026). doi:10.1126/sciimmunol.aec5171

75 Feau, S., Arens, R., Togher, S. & Schoenberger, S. P. Autocrine IL-2 is required for secondary population expansion of CD8+ memory T cells. Nature Immunology 12, 908–913 (2011). 10.1038/ni.2079

76 Bosteels, V. & Janssens, S. Striking a balance: new perspectives on homeostatic dendritic cell maturation. Nat Rev Immunol 25, 125–140 (2025). 10.1038/s41577-024-01079-5

77 Tur, C. et al. CD19-CAR T-cell therapy induces deep tissue depletion of B cells. Ann Rheum Dis 84, 106–114 (2025). 10.1136/ard-2024-226142

78 Bhoj, V. G. et al. Persistence of long-lived plasma cells and humoral immunity in individuals responding to CD19-directed CAR T-cell therapy. Blood 128, 360–370 (2016). 10.1182/blood-2016-01-694356

79 Tu, E. et al. T Cell Receptor-Regulated TGF-beta Type I Receptor Expression Determines T Cell Quiescence and Activation. Immunity 48, 745–759 e746 (2018). 10.1016/j.immuni.2018.03.025

80 Ogishi, M. et al. Impaired development of memory B cells and antibody responses in humans and mice deficient in PD-1 signaling. Immunity 57, 2790–2807 e2715 (2024). 10.1016/j.immuni.2024.10.014

81 Love, M. I., Huber, W. & Anders, S. Moderated estimation of fold change and dispersion for RNA-seq data with DESeq2. Genome Biol 15, 550 (2014). 10.1186/s13059-014-0550-8

82 Racle, J. & Gfeller, D. in Bioinformatics for Cancer Immunotherapy: Methods and Protocols (ed Sebastian Boegel) 233–248 (Springer US, 2020).

83 Bergen, V., Lange, M., Peidli, S., Wolf, F. A. & Theis, F. J. Generalizing RNA velocity to transient cell states through dynamical modeling. Nat Biotechnol 38, 1408–1414 (2020). 10.1038/s41587-020-0591-3

